# Antisense transposon insertions into host genes trigger piRNA mediated immunity

**DOI:** 10.1101/2025.07.28.667215

**Authors:** Baptiste Rafanel, Liudmila Protsenko, Dominik Handler, Julius Brennecke, Kirsten-André Senti

## Abstract

Transposable elements pose a persistent threat to genome integrity, yet how host defense systems adapt to newly invading elements remains poorly understood. Here, we reveal how *Drosophila melanogaster* acquired PIWI-interacting RNA (piRNA)-mediated immunity against the recently invading endogenous retrovirus *tirant*. By integrating genetics, small RNA profiling, and population genomics, we identify two distinct modes of de novo piRNA biogenesis. The primary mechanism involves antisense insertions into the *flamenco* cluster, a well-established master locus for transposon control. Strikingly, we also find that antisense *tirant* insertions into 3′ UTRs of host genes robustly trigger piRNA production, a process driven by host gene transcription but independent of gene identity. These findings challenge prevailing models that link piRNA precursor specification to genomic origin or nuclear processing context. Instead, they uncover a flexible, general mechanism in which transposition into host gene exons represents a critical vulnerability for transposons: by generating chimeric antisense transcripts that are exported to the cytoplasm, transposons inadvertently initiate their own silencing, enabling rapid and adaptive genome defense against new invaders.

## INTRODUCTION

The PIWI-interacting RNA (piRNA) pathway safeguards germline genome integrity in animals by silencing transposable elements. In this adaptive, small RNA-based defense system, 22–32 nucleotide piRNAs guide PIWI-clade Argonaute proteins to complementary transposon transcripts, directing transcriptional and post-transcriptional silencing mechanisms (Siomi *et al*, 2011; Czech *et al*, 2018; Ozata *et al*, 2019). piRNAs are generated in the cytoplasm from single-stranded transcripts via two interconnected processes: phased biogenesis, in which precursors are cleaved stepwise in a 5′ to 3′ direction (Han *et al*, 2015; Mohn *et al*, 2015; Homolka *et al*, 2015), and the ping-pong amplification cycle, which enhances piRNA abundance through reciprocal cleavage of complementary sense and antisense transcripts (Brennecke *et al*, 2007; Gunawardane *et al*, 2007). Together, these mechanisms constitute a conserved molecular framework for piRNA biogenesis across metazoans (Gainetdinov *et al*, 2018).

piRNAs predominantly derive from discrete genomic loci known as piRNA clusters. These loci are often enriched in fragmented transposon insertions and are thought to function as molecular memory banks of past transposon invasions (Aravin *et al*, 2007; Brennecke *et al*., 2007; Houwing *et al*, 2007; Malone *et al*, 2009). In *Drosophila*, for example, the *flamenco* and *77B* clusters are essential for silencing specific transposons (Pelisson *et al*, 1994; Sarot *et al*, 2004; Brennecke *et al*., 2007; Senti *et al*, 2025), while others may target inactive elements or act redundantly (Gebert *et al*, 2021). Collectively, these observations have led to the "trap model" of piRNA biogenesis, which proposes that a transposon becomes targetable only after inserting into an active piRNA cluster and thereby donating its sequence to the system (Bergman *et al*, 2006; Brennecke *et al*., 2007; Zanni *et al*, 2013).

Yet this model raises a conceptual problem. Like most cellular RNAs, piRNA precursors are transcribed by RNA polymerase II as single-stranded transcripts (Senti & Brennecke, 2010). What, then, distinguishes an RNA destined for piRNA processing from a conventional mRNA? Current models refer to various criteria, including specific RNA motifs or structures (Homolka *et al*., 2015; Ishizu *et al*, 2015), chromatin context or nuclear processing events unique to piRNA precursors (Le Thomas *et al*, 2014b; Zhang *et al*, 2014; Yu *et al*, 2019), and piRNA-guided slicing (Han *et al*., 2015; Mohn *et al*., 2015; Homolka *et al*., 2015). However, the underlying molecular rules that define a piRNA precursor remain largely unclear. Further complicating matters, the genetic and epigenetic features that specify piRNA cluster identity vary across species and cell types (Malone *et al*., 2009; Robine *et al*, 2009; Li *et al*, 2013; Mohn *et al*, 2014), and isolated transposon insertions outside of clusters can also elicit piRNA production (Shpiz *et al*, 2014; Mohn *et al*., 2014; Baumgartner *et al*, 2022), challenging the universality of the trap model.

Dissecting how the piRNA pathway distinguishes its precursors from other cellular RNAs is challenging in established transposon control settings, where the initial triggers for piRNA production are obscured by layers of evolutionary adaptation. In the *Drosophila* germline, one of the most powerful study systems for piRNA biology, piRNA clusters are epigenetically defined and rely on the Rhino-Deadlock-Cutoff (RDC) complex to generate piRNA precursor transcripts (Josse *et al*, 2007; Brennecke *et al*, 2008; Klattenhoff *et al*, 2009; Le Thomas *et al*, 2014a; Mohn *et al*., 2014; Zhang *et al*., 2014). However, this chromatin-based mechanism is not conserved outside of fruit flies, and the interplay between maternally inherited piRNAs and chromatin context further complicates the identification of general rules underlying piRNA precursor specification.

To address this, we turned to the simplified piRNA pathway that operates in somatic cells of the *Drosophila* ovary. This pathway targets insect endogenous retroviruses (iERVs) of the *Ty3/gypsy* class, which are specifically expressed in somatic follicle cells to form enveloped viral particles that infect the neighboring germline (Pelisson *et al*., 1994; Leblanc *et al*, 2000; Sarot *et al*., 2004; Senti *et al*., 2025). Importantly, the somatic piRNA pathway lacks the RDC system, operates independently of maternally inherited piRNAs, and is therefore free from germline-specific chromatin regulation (Senti & Brennecke, 2010; Czech *et al*., 2018).

By leveraging natural *Drosophila melanogaster* populations that have recently experienced the horizontal transfer of an active transposon, we set out to dissect how naïve genomes initiate somatic piRNA production de novo, unconfounded by pre-existing immunity or laboratory conditions. Although once thought rare, at least ten such invasions have occurred in *Drosophila melanogaster* over the past two centuries (Bartolome *et al*, 2009; Scarpa *et al*, 2024; Pianezza *et al*, 2025; Scarpa *et al*, 2025). One such transposon is the iERV *tirant*, which likely invaded *D. melanogaster* from *D. simulans* in the mid- 20th century (Akkouche *et al*, 2012; Schwarz *et al*, 2021). Since then, *tirant* has spread worldwide, offering a unique opportunity to dissect the molecular events that trigger piRNA biogenesis in response to a novel genome invader.

Here, we show that natural *D. melanogaster* populations have evolved piRNA-mediated resistance to *tirant* via two distinct mechanisms. As predicted by the trap model, single antisense insertions of *tirant* into the *flamenco* piRNA cluster are sufficient to initiate piRNA production and silence the element. Strikingly, we also uncovered a second, equally potent mechanism: antisense *tirant* insertions into the 3′ UTRs of host genes trigger robust piRNA production and transposon repression. We show that piRNA biogenesis from these insertions requires host gene transcription yet is independent of gene identity. These findings challenge the view that piRNA precursor identity is defined by genomic origin or nuclear RNA processing history. Instead, they highlight the importance of antisense transposon sequences as part of chimeric host transcripts as a trigger for de novo piRNA biogenesis. Our results provide direct support for a model proposed in vertebrates and mosquitoes, in which antisense insertions near host gene 3′ ends can serve as entry points for piRNA immunity against endogenous retroviruses, exogenous RNA viruses, and transposable elements (Konstantinidou *et al*, 2024; Qu *et al*, 2023; Yu *et al*, 2025). Together, this work defines a generalizable framework for how small RNA-based immunity evolves in response to transposon invasions.

## RESULTS

### Natural *D. melanogaster* populations evolved diverse piRNA responses against *tirant*

*tirant* belongs to the *ZAM* subclade of *Ty3/gypsy*-class iERVs and encodes three open reading frames: *gag*, *pol*, and *env* (Figure 1A, B) (Marsano *et al*, 2000; Akkouche *et al*., 2012; Schwarz *et al*., 2021; Senti *et al*., 2025). To investigate how natural *D. melanogaster* populations responded to the reported *tirant* invasion during the 20th century (Schwarz *et al*., 2021), we analyzed the founder lines of the Drosophila Synthetic Population Resource (DSPR), a panel of fifteen strains collected from diverse geographic regions between the years 1930 and 1970 (Figure 1C) (King *et al*, 2012). High-quality, long-read genome assemblies are available for all available DSPR strains except *B7* (Chakraborty *et al*, 2018; Chakraborty *et al*, 2019). Because the original *B1* and *AB8* strains are no longer accessible, we used the *Ber2* and *Sam* strains as proxies as they were collected at the same respective locations and time points. In addition to the DSPR panel, we analyzed the *iso-1* strain, a mosaic line derived from laboratory stocks of uncertain origin and collection date (Brizuela *et al*, 1994). This strain forms the basis of the current *D. melanogaster* reference genome (Hoskins *et al*, 2015).

**Figure 1.**
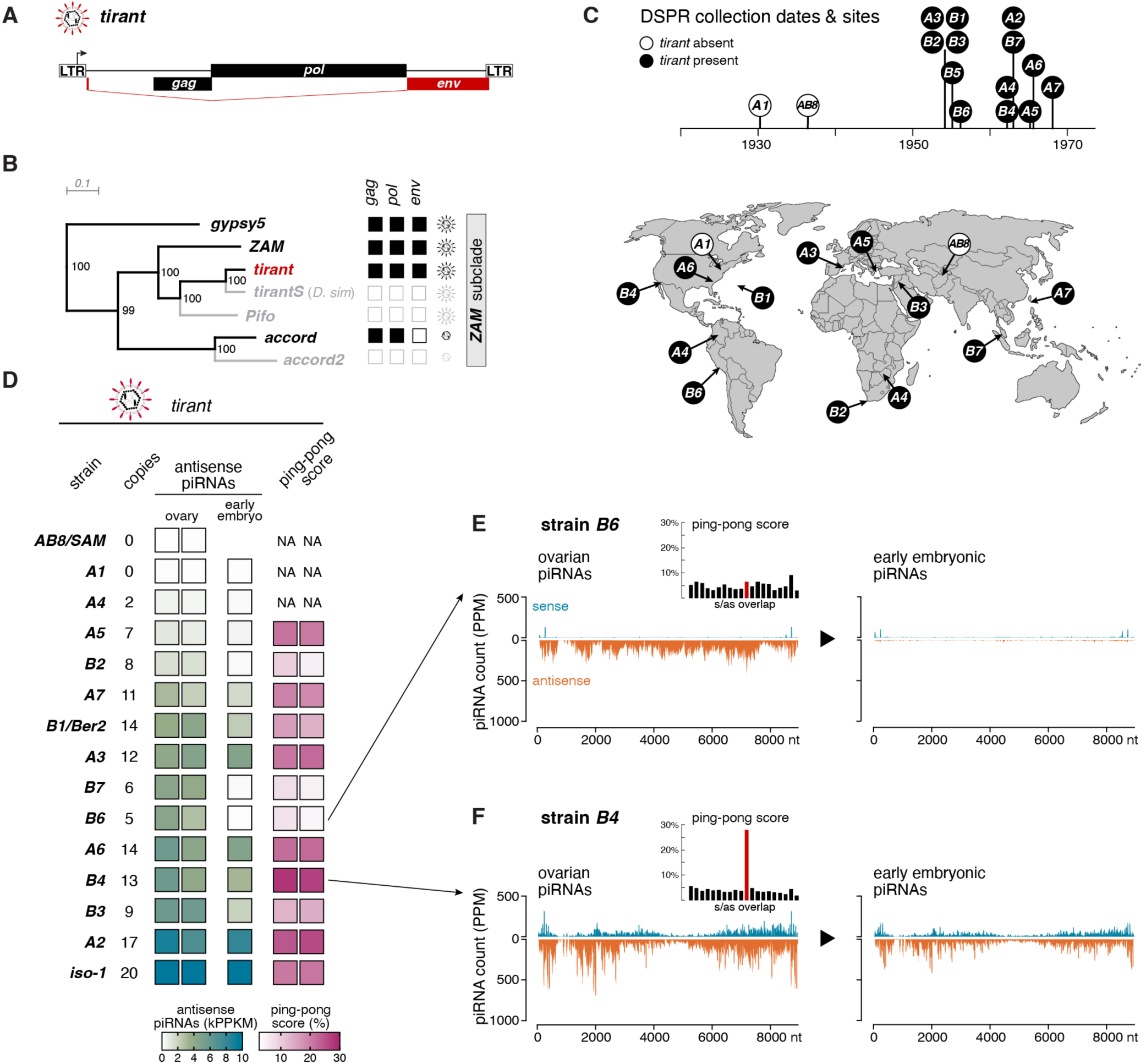
Natural *D. melanogaster* populations evolved diverse piRNA responses against *tirant*. (A) Schematic of the *tirant* retrotransposon showing LTRs and ORFs (*gag, pol, env*). (B) Phylogenetic tree of the *ZAM* subclade of the *gypsy/gypsy* clade iERVs based on full-length Pol sequences. Right: ORF integrity (black: intact; white: broken). Scale bar: amino acid substitutions/site. (C) Timeline (top) and world map (bottom) of DSPR founder strain collections. *A1* and *AB8/Sam* (white) lack *tirant*; other strains (black) contain *tirant* copies (map adapted from Chakraborty et al., 2019; www.outline-world-map.com). (D) Table (left) showing estimated *tirant* copy numbers from genomic Illumina data and heatmaps (right) of 23–29 nt antisense piRNA levels (thousand reads per kb, normalized to 1M miRNAs) mapping perfectly to *tirant* in ovaries and 0–30 min embryos as well as ping-pong scores for ovary samples (NA: insufficient piRNAs for calculation in *A1*, *A4*, *AB8/Sam*). Ovarian small RNAs determined in biological duplicate. (E) Density plots of sense (positive) and antisense (negative) *tirant*-mapping piRNAs (23–29 nt, PPM) along the consensus *tirant* sequence in *B6* ovaries (left) and early embryos (right). Inset: 5′ overlap histogram of ovarian piRNAs (10 nt ping-pong overlap in red). (F) As in (E), for strain *B4*.

We first determined the presence of *tirant* in each strain by comparing their genomes to the *tirant* consensus sequence. Two strains (*A1/Canton-S* and *AB8/Sam*) lacked *tirant* entirely, whereas the remaining twelve harbored variable copy numbers (Figure 1D), often including at least one structurally intact insertion, indicative of ongoing or recent activity. The collection dates of the DSPR strains align with prior work that placed the *tirant* invasion between 1940 and 1950 (Schwarz *et al*., 2021). Notably, *iso-1* carries the highest number of *tirant* insertions among all strains analyzed (Kaminker *et al*, 2002).

To test whether strains with *tirant* insertions established corresponding piRNA responses, we sequenced Argonaute-bound small RNAs from dissected ovaries and early embryos (Grentzinger *et al*, 2020). Ovarian small RNAs encompass both germline and somatic piRNAs, while small RNAs from early embryos serve as a readout of maternally deposited germline piRNAs (Malone *et al*., 2009). All *tirant* positive strains, including *iso-1*, expressed *tirant* piRNAs in ovaries, although their abundance and biogenesis features varied substantially (Figure 1D).

Two distinct piRNA pathways operate in the *Drosophila* ovary: one in germline cells and one in the surrounding somatic follicle cells (Vagin *et al*, 2006; Pelisson *et al*, 2006; Brennecke *et al*., 2007; Gunawardane *et al*., 2007; Klattenhoff *et al*., 2009; Malone *et al*., 2009; Han *et al*., 2015; Mohn *et al*., 2014). Germline cells express Piwi, Aubergine, and Ago3, which together mediate phased and ping-pong piRNA biogenesis. This results in mixed populations of sense and antisense piRNAs, enriched for a characteristic 10-nt overlap, known as the ping-pong signature (Figure S1A-C; illustrated with *burdock*, a germline-targeted transposon). In contrast, somatic follicle cells express only Piwi and generate piRNAs exclusively via phased processing from long, single-stranded precursor RNAs. These somatic piRNAs are typically antisense and lack a ping-pong signature (Figure S1D-F; illustrated with *gypsy5*, a soma-targeted transposon).

Analysis of *tirant* piRNAs revealed that most strains produce both sense and antisense piRNAs, with varying degrees of ping-pong amplification (Figure 1D-F). Such profiles are consistent with piRNA production in the germline alone, or in both germline and soma. However, some strains, such as *B6* and *B7*, produced exclusively antisense piRNAs without ping-pong features, indicative of a strictly somatic origin. Supporting this, these strains also lacked maternally inherited *tirant* piRNAs in early embryos (Figure 1E).

Altogether, these data reveal that natural *D. melanogaster* populations evolved piRNA responses against the recently invading *tirant* iERV. However, the variability in the piRNA patterns among fly strains raised the question of which cell types express *tirant*, and whether both the germline and somatic piRNA pathways contribute to its silencing.

### The somatic piRNA pathway silences *tirant*

To determine the site of *tirant* expression in the ovary, we generated transcriptional reporters in which the *tirant cis*-regulatory sequences (LTR and 5′ UTR) drove either β-Galactosidase (*tirant*-lacZ) or a GFP–β-Galactosidase fusion (*tirant*-GFP:lacZ) (Figure 2A). Each construct was integrated into an *attP* landing site on the second chromosome of a laboratory strain devoid of *tirant* piRNAs (Figure 2B). In this piRNA-naïve background, both reporters were strongly and selectively expressed in the ovarian soma, with no detectable germline activity (Figure 2C, D). However, crossing these reporters to the *iso- 1* strain, which produces abundant *tirant* piRNAs, resulted in silencing, reducing reporter activity to undetectable levels (Figure 2E–G).

**Figure 2:**
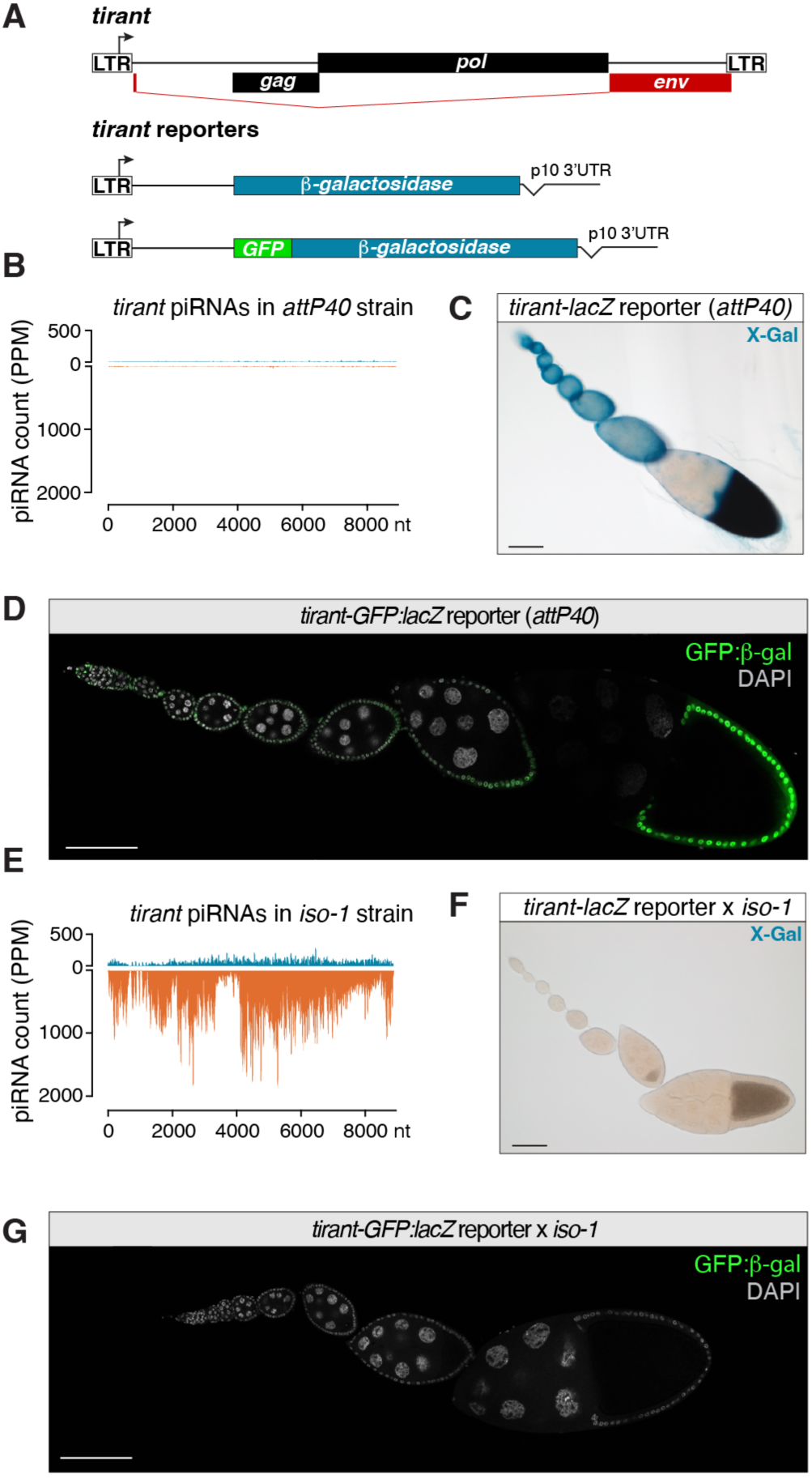
A *tirant* reporter is expressed in follicle cells and silenced by piRNAs. (A) Schematic depicting the design of the *tirant-lacZ and GFP:lacZ* reporters. (B) Density plot of *tirant*-mapping piRNAs (PPM) along the *tirant* consensus sequence in the *attP40* strain. (C) X-gal staining of an ovariole of transgenic *tirant- lacZ* reporter flies in the permissive background (*attP40*). Scale bar: 100μm. (D) GFP signal of the *tirant-GFP:lacZ* reporter in an ovariole in the permissive genetic background (*attP40*). DNA staining (DAPI) is shown in grey. Scale bar: 100μm. (E) Density plot of *tirant*-mapping piRNAs (PPM) along the *tirant* consensus sequence in the *iso-1* strain. (F) As in (D) but showing an ovariole from the progeny of the cross of transgenic *tirant-lacZ* reporter females to *iso-1* males. (G) As in (C). but showing an ovariole from the progeny of the cross of transgenic *tirant-GFP:lacZ* reporter females to *iso-1* males.

To investigate the expression of endogenous *tirant* retroviruses, we disrupted the piRNA pathway in flies carrying full-length *tirant* insertions using transgenic RNA interference (RNAi) (Dietzl *et al*, 2007; Ni *et al*, 2011; Handler *et al*, 2013). This strategy was infeasible in standard laboratory strains as fly lines for tissue-specific RNAi either lacked *tirant* entirely (*tj*-Gal4 crossed to VDRC dsRNA lines targeting the somatic pathway) or harbored only fragmented insertions (*MTD-Gal4* crossed to short TRiP hairpin lines for germline knockdown) (Figure S2). To overcome these limitations, we generated introgressed lines by crossing *iso-1*, harboring multiple full-length *tirant* insertions across all major chromosomes, into selected ovarian Gal4 drivers and UAS-RNAi strains. This approach allowed us to interrogate *tirant* expression in its native genomic context under defined pathway perturbations.

Using RNA fluorescence in situ hybridization (RNA-FISH), we detected no *tirant* expression in strains lacking insertions (e.g., *A1*) or in unmanipulated *iso-1*, despite harboring ∼ 20 full-length copies. In contrast, disruptiom of the somatic piRNA pathway in the introgressed background resulted in strong *tirant* expression across the ovarian soma. Transcripts accumulated in the nuclei and at the apical membranes of follicle cells, with clear signal also observed in late-stage oocytes, indicating effective soma-to-germline transmission of *tirant* RNA (Figure 3A) as previously reported for the related iERV *ZAM* (Yoth *et al*, 2023; Senti *et al*., 2025). By contrast, germline piRNA pathway knockdown did not lead to *tirant* derepression in either germline or soma (Figure 3B). Consistent with these observations, small RNA sequencing of ovaries depleted for germline piRNAs (*MTD-Gal4*-driven *aub*/*ago3* knockdown) revealed persistant and abundantly produced *tirant* piRNAs bearing hallmarks of somatic origin, including a pronounced antisense bias and the absence of a ping-pong signature (Figure 3C; ping-pong Z-score: 0.8).

**Figure 3:**
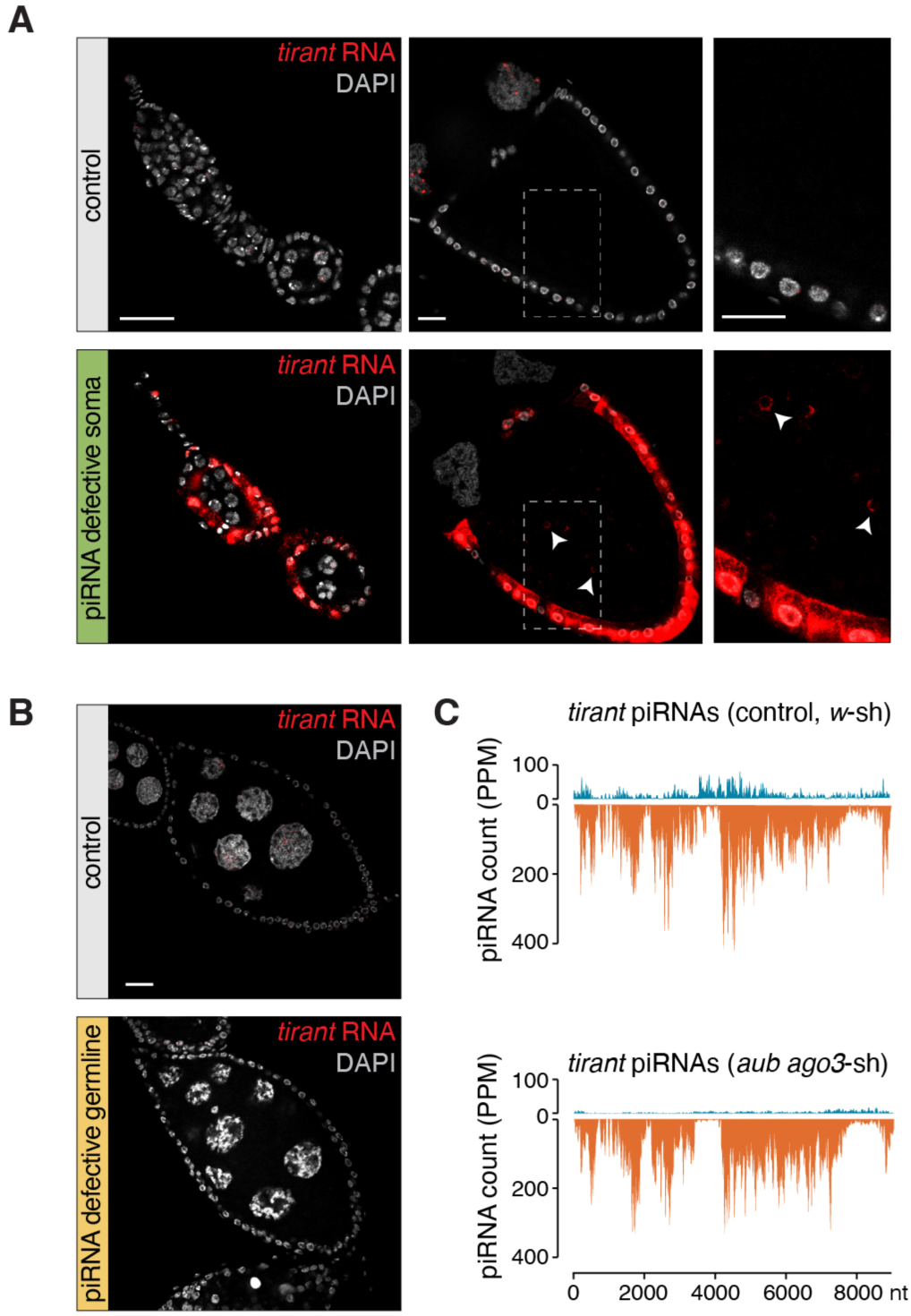
The somatic piRNA pathway silences the *tirant* retrovirus. (A) RNA-FISH detecting *tirant* sense transcripts (red) in early and late-stage egg chambers of control ovaries (*arr2* knockdown; top) or ovaries with defective somatic piRNA pathway (*vret* knockdown; bottom). Both genotypes harbor active *tirant* copies. Right panels are zoom-in from the boxed parts of the middle panels. Arrowheads show *tirant* RNA detected in the oocyte. DNA staining (DAPI) is shown in grey. Scale bar: 20μm. (B) RNA-FISH detecting *tirant* sense transcripts (red) in stage 8 egg chambers with a germline specific RNAi knockdown of a control gene (*white;* top) or of the germline piRNA pathway (*aub* and *ago3*; bottom). Both genotypes harbor active *tirant* copies. DNA staining (DAPI) is shown in grey. Scale bar: 20μm. (C) Density plot of *tirant*-mapping piRNAs (PPM) along the *tirant* consensus sequence in the strains shown in (B).

Altogether, our findings demonstrate that the *tirant* iERV is intrinsically transcribed in the ovarian soma but efficiently silenced by the somatic piRNA pathway. Consistent with its ability to infect the germline, *tirant* encodes an intact Env-F glycoprotein like other somatically expressed, infectious iERVs (Pelisson *et al*., 1994; Leblanc *et al*, 2000; Marsano *et al*., 2000; Senti *et al*., 2025).

### Independent antisense insertions in *flamenco* underlie *tirant* silencing in most natural strains

The DSPR founder panel provided a powerful framework to investigate how natural *D. melanogaster* populations evolved resistance to *tirant*. These strains represent diverse geographic origins, were largely collected after the estimated *tirant* invasion, and possess high-quality genome assemblies that allow the accurate mapping of transposon insertions, even in complex heterochromatic regions (Chakraborty *et al*., 2019; Chakraborty *et al*., 2018; King *et al*., 2012).

Given that *tirant* is transcribed in the ovarian soma and silenced by the somatic piRNA pathway (Figure 3), we asked whether all natural *tirant*-carrying strains produce functional *tirant*-targeting piRNAs in this tissue. To test this, we crossed each strain to the *tirant-lacZ* reporter line, which lacks *tirant*-derived piRNAs (Figure 2B). Reporter silencing in follicle cells served as a direct functional readout for somatic piRNA-mediated repression (Sarot *et al*., 2004).

As expected, *tirant*-naïve strains such as *A1* and *AB8*/*Sam* failed to silence the reporter (Figure 4A, B). In contrast, all *tirant*-positive strains repressed the reporter to varying degrees: eight showed complete repression (e.g., *B2*), while five displayed partial (e.g., *A6*) or weak (e.g., *A4*) repression (Figure 4A–C; Figure S3). Reporter repression occurred independent of the parental crossing direction, ruling out a confounding role for maternally inherited piRNAs.

**Figure 4:**
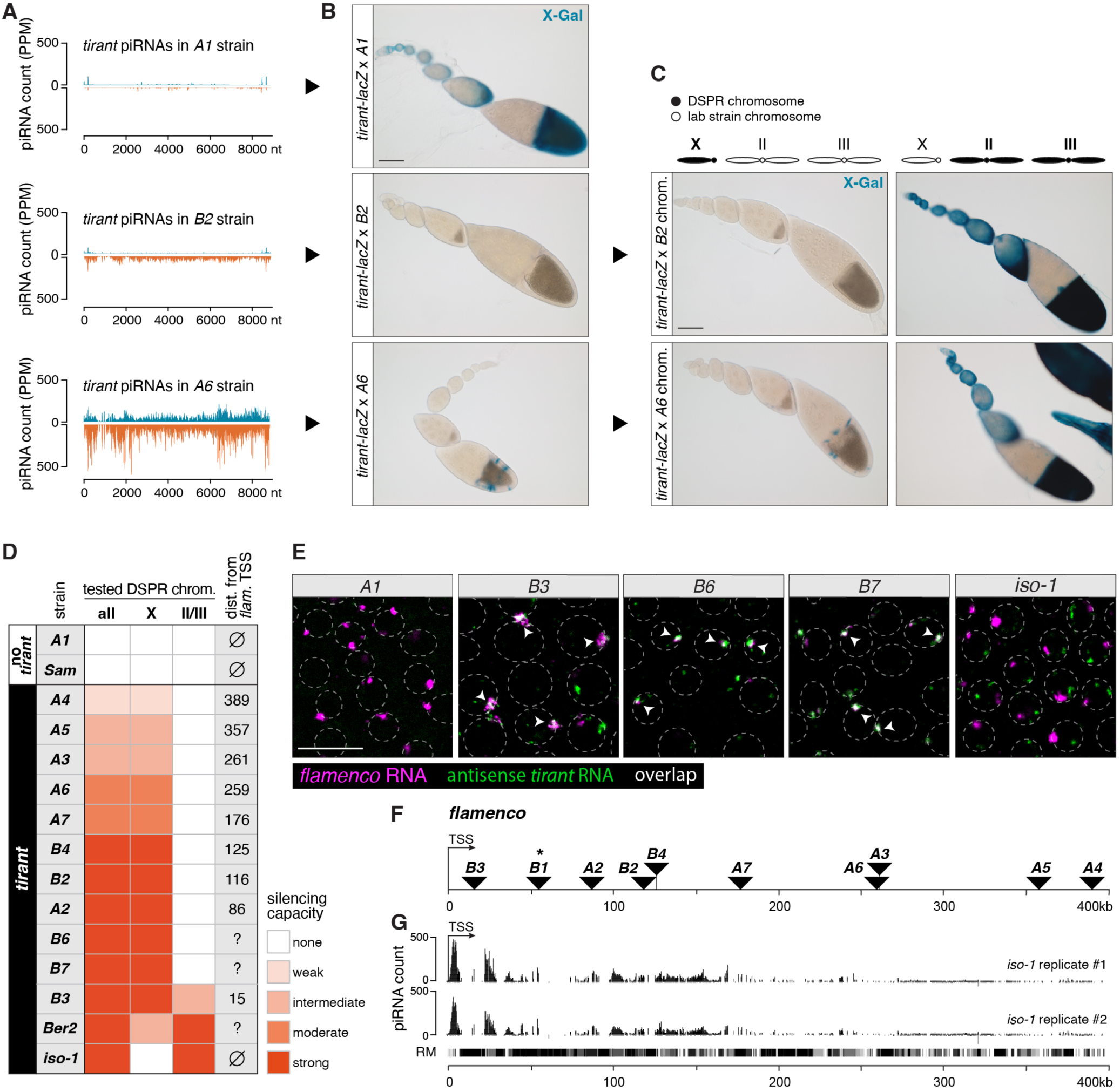
Independent antisense insertions in *flamenco* underlie *tirant* silencing in most natural strains. (A) Density plot of piRNAs (PPM) mapping to the *tirant* consensus sequence from the *A1* strain (top), the *B2* strain (middle), or the *A6* strain (bottom). (B) X-gal stainings of ovarioles from the progeny of crosses of transgenic *tirant-lacZ* reporter females to males from DSPR founder strains shown in (A). Scale bar: 100μm. (C) X-gal stainings of ovarioles from the progeny of crosses between transgenic *tirant-lacZ* reporter females to males from hybrid strains harboring the indicated DSPR founder chromosomes of *B2* (top) or *A6* (bottom). Left panels show the tested X chromosomes from DSPR strains, right panels the autosomes from tested DSPR strains. Scale bar: 100μm. (D) Summary of the *tirant-lacZ* reporter silencing capacity of the different DSPR strains (all) and the contribution of the X chromosome or the pair of autosomes for each DSPR strain. The distance between the transcriptional start site (TSS) of *flamenco* and the *tirant* insertion is shown to the right (distance could not be calculated for the *B6*, *B7*, and *Ber2* strains). (E) RNA-FISH detecting *flamenco (*magenta) and antisense *tirant* (green) transcripts in somatic follicle cell nuclei of stage 8 egg chambers of indicated DSPR strains. Somatic follicle cell nuclei circumferences are marked by dashed lines (inferred from DAPI staining). Colocalization of magenta and green signal results in white (marked by arrow heads). Scale bar: 20μm. (F) Plot showing the localization of *tirant* insertions (black triangles) in *flamenco* for each DSPR strain relative to the *flamenco* transcriptional start site (TSS). * indicates that the contribution of the *tirant flamenco* insertion in *B1* could not be directly assessed. (G) UCSC genome browser screenshot of the *flamenco* locus in *iso-1* (400kb downstream of the *flamenco* TSS) with ovarian genome-unique mapping piRNAs (PPM) from the *iso-1* strain shown. The repeat masker (RM) track is shown at the bottom.

To map the genomic sources of the *tirant*-repressive piRNAs, we generated hybrid lines in which either the X chromosome or the major autosomes (chromosomes 2 and 3) from each *tirant*-positive strain were introduced into a balancer background devoid of *tirant* piRNAs. Crossing these hybrids to the *tirant-lacZ* reporter revealed the chromosomal origin of functional piRNA source loci. For example, in hybrids from strains *B2* and *A6*, repression occurred only when the DSPR-derived X chromosome was present, pinpointing the piRNA source to the X chromosome (Figure 4B–D; Figure S3).

Notably, in ten of the thirteen repressive strains, functional piRNA source activity mapped exclusively to the X chromosome (Figure 4C). To identify these loci, we examined all X-linked *tirant* insertions and analyzed piRNA production from their flanking sequences as a proxy for source activity. Of 45 insertions, six stand-alone euchromatic copies produced weak, bidirectional piRNAs typical of germline piRNA source loci (Figure S4, Table S1) (Shpiz *et al*., 2014; Mohn *et al*., 2014). Only one euchromatic insertion (in *A3*) produced low-level unistranded piRNAs suggestive of a weak somatic source locus (see below).

In contrast, eight of the ten repressive strains harbored antisense *tirant* insertions within *flamenco*, the major somatic piRNA cluster located near the pericentromeric heterochromatin of the X chromosome (Brennecke *et al*., 2007; Pelisson *et al*., 1994; Sarot *et al*., 2004; Senti *et al*., 2025). For the remaining two strains (*B6* and *B7*) a direct assessment was not possible: *B6* lacks a fully assembled *flamenco* locus, and *B7* has no sufficiently assembled genome due to missing long-read data. However, in both cases, RNA-FISH revealed co-localization of *tirant* antisense transcripts with *flamenco* sense RNA in follicle cell nuclei, mirroring strain *B3* where a *tirant* insertion in *flamenco* is confirmed (Figure 4E). These observations strongly suggest that *B6* and *B7* also carry *tirant* insertions in *flamenco*. Notably, with the exception of *A3*, *A6*, and *A7* (likely representing a shared insertion), the *tirant* insertions in *flamenco* appear to have occurred independently across strains.

*flamenco* is a unistrand piRNA cluster transcribed from a single promoter and spanning several hundred kilobases (Brennecke *et al*., 2007; Goriaux *et al*, 2014; Mohn *et al*., 2014; Senti *et al*., 2025). Although all identified *tirant* insertions within *flamenco* were single-copy, the silencing efficacy varied widely across strains, ranging from complete repression to minimal effect (Figure 4C). This variability correlated with insertion position relative to the *flamenco* promoter: proximal insertions conferred strong silencing, whereas distal insertions were much less effective (Figure 4C,F). For instance, strain *A2*, with a *tirant* insertion 86 kb from the promoter, fully repressed the reporter, while strain *A4*, harboring an insertion ∼400 kb downstream, showed minimal repression. Interestingly, this trend aligns with a gradient of piRNA output across *flamenco*, which declines with increasing distance from the promoter, underscoring the positional impact of integration site on silencing efficacy (Figure 4G; Table S1).

During this analysis, we also identified multiple RepeatMasker-annotated degenerate *tirant*-like fragments within *flamenco*. These correspond to *pifo*, a non-functional *tirant*-related element largely confined to *flamenco* (Zanni *et al*., 2013). Since *pifo* fragments were present in all strains, including those incapable of repressing the *tirant* reporter, we conclude that *pifo*-derived piRNAs, which share ∼60% sequence identity with *tirant*, do not effectively cross-silence *tirant*.

Together, these findings demonstrate that in most natural strains, *tirant* repression evolved through independent antisense insertions into *flamenco*, confirming *flamenco* as the primary source of somatic piRNAs. This mirrors observations in *D. simulans*, where the recently invading iERV *Shellder* similarly acquired independent antisense insertions into *flamenco* across multiple strains (Scarpa *et al*., 2025).

### Antisense insertions into host gene 3′ UTRs provide an alternative silencing route

In three strains, *iso-1*, *B3*, and *Ber2*, *tirant* silencing mapped partly or entirely to the autosomes (Figure 4C). In *Ber2*, autosomal activity exceeded that of the X chromosome, while in *iso-1*, silencing was exclusively autosomal. To uncover the origin of these non-*flamenco* piRNA sources, we first focused on *iso-1*, where silencing mapped specifically to chromosome 2, a region lacking any previously described somatic piRNA clusters (Figure 5A).

**Figure 5:**
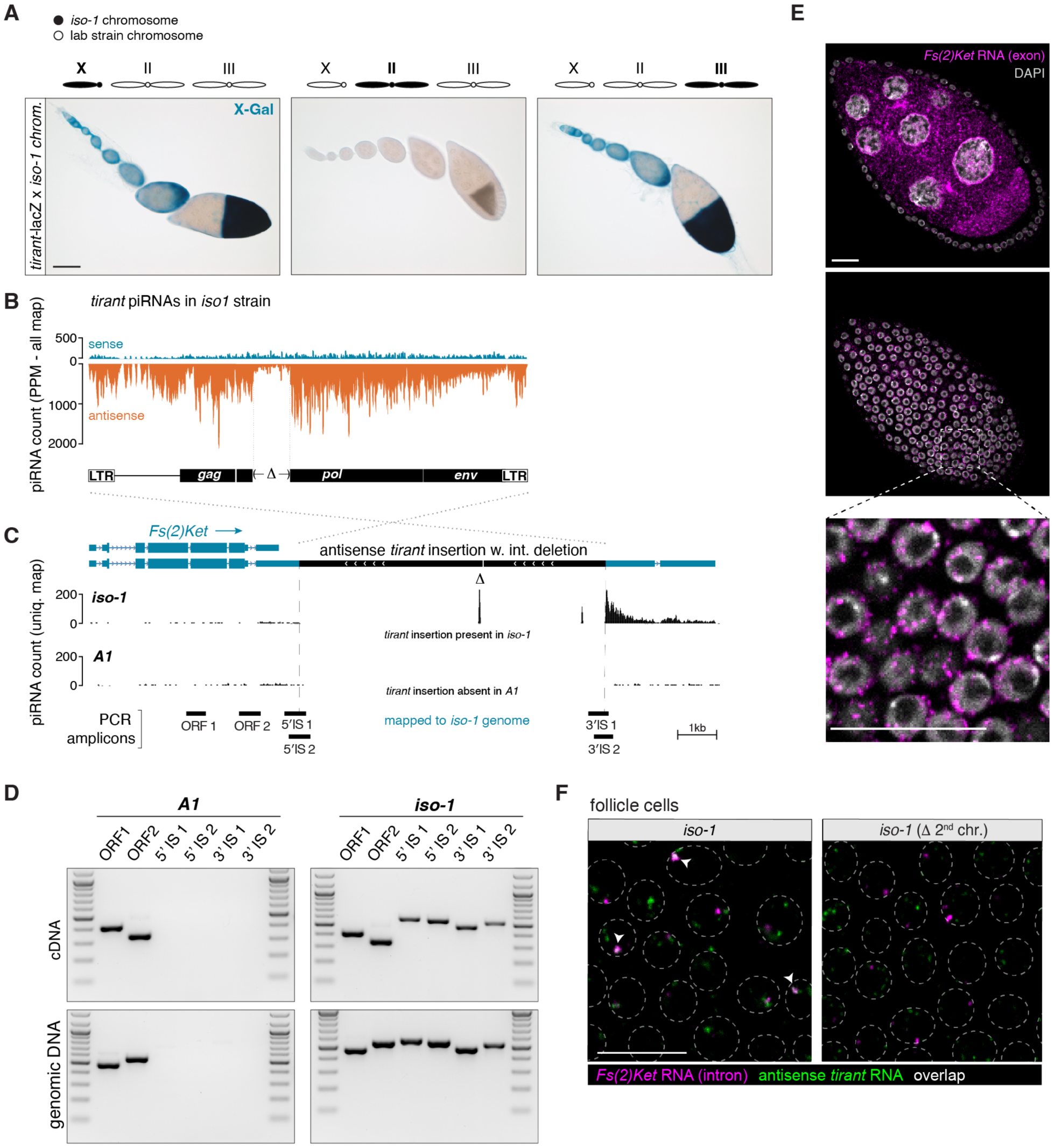
Antisense insertions into host gene 3′ UTRs provide an alternative silencing route. (A) X-gal stainings of ovarioles from the progeny of crosses between transgenic *tirant-lacZ* reporter females and males from strains harboring either the X (left), 2^nd^ (middle), or 3^rd^ chromosome (right) from *iso-1* as indicated. Scale bar: 100μm. (B) Density plot of piRNAs (PPM) from the *iso-1* strain mapping to the *tirant* consensus sequence. Dashed lines mark the region with distinctly lower piRNA coverage, corresponding to the deletion in the *tirant* insertion in *Fs(2)Ket*. (C) UCSC browser screenshot of the *Fs(2)Ket* locus in the *iso-1* genome. The two tracks show genome-unique piRNAs (in PPM) sequenced from ovaries of *iso-1* (top) or *A1* (bottom). piRNAs mapping uniquely to the internal deletion of *tirant* are labeled by Δ. Positions of analytic PCR amplicons shown in (E) are indicated at the bottom. Scale bar: 1kb. (D) RT-PCR on RNA (top) and PCR on genomic DNA (bottom) from the *A1* and *iso-1* strains to detect chimeric transcripts between *Fs(2)Ket* and *tirant*. Positions of PCR amplicons are shown in panel C. Marker: 100bp DNA ladder. (E) RNA-FISH detecting *Fs(2)Ket* transcripts (exonic probes) in germline (top), and somatic cells (middle) with a zoom-in from the boxed part shown below. Images correspond to a stage 7/8 egg chamber isolated from the *A1* strain. Scale bar: 20μm. (F) RNA-FISH detecting intronic *Fs(2)Ket* (magenta) and antisense *tirant* (green) transcripts in follicle cells from stage 8 egg chambers from *iso-1* (left) and in an *iso-1* strain where the second chromosome (containing the *Fs(2)Ket* locus) was replaced by balancer chromosomes (*iso-1(Δ2^nd^ chr.);* right). Circumferences of follicle cell nuclei are marked by dashed lines. Arrowheads indicate foci containing both signals. Scale bar: 20μm.

Among the *tirant* insertions on chromosome 2, one stood out: it contained a 752 bp internal deletion coinciding with a sharp drop in antisense piRNA coverage across the element (Figure 5B). This insertion resides in antisense orientation within the long 3′ UTR isoform of *Fs(2)Ketel (Fs(2)Ket)*, a ubiquitously expressed nuclear importin-β gene (Figure 5C) (Lippai *et al*, 2000). Consistent with this locus acting as a piRNA source, we detected abundant piRNAs in *iso-1*, but not in other strains such as *A1*, mapping to the deletion junction and the downstream flanking region indicative of phased piRNA production (Figure 5C). The lack of 3′ UTR piRNAs in the *A1* strain indicates that the production of piRNAs in *iso-1* is not an intrinsic feature of the *Fs(2)Ket* 3′ UTR.

RT-PCR on total ovarian RNA using junction-spanning amplicons confirmed that the *tirant* insertion is transcribed as part of the *Fs(2)Ket* 3′ UTR (Figure 5C, D). Since *Fs(2)Ket* is expressed in both soma and germline (Figure 5E), we used RNA FISH with intronic probes for *Fs(2)Ket* to determine whether the *tirant* insertion is transcribed in somatic follicle cells. Indeed, antisense *tirant* transcripts co-localized with the *Fs(2)Ket* gene locus in follicle cell nuclei but not with *flamenco* (Figure 5F, 3E), supporting expression as a chimeric host–transposon transcript.

To assess whether *Fs(2)Ket* in *iso-1* was the sole autosomal *tirant* piRNA source locus, we examined *Ber2* (a proxy for *B1*) and *B3*. Remarkably, *Ber2* also harbors an antisense *tirant* insertion in the *Fs(2)Ket* 3′ UTR, producing unistranded piRNAs that extend downstream (Figure S5). As in *iso-1*, RT-PCR and RNA-FISH confirmed host gene-driven transcription of the insertion (Figure S5). Despite occupying the exact same nucleotide position (verified by PCR-sequencing in *Ber2*; Figure S5F), the *iso-1* and *Ber2* insertions are independent events, as revealed by distinct SNP profiles reflecting strain-specific *tirant* polymorphisms (Figure S5G).

In *B3*, we identified a third autosomal piRNA source: an antisense *tirant* insertion in the 3′ UTR of an alternative *cactus* transcript isoform. No other chromosome 2 insertions in *B3* produced detectable piRNAs. Although piRNA levels from this locus were modest, consistent with *B3*’s relatively weak autosomal silencing, the piRNA reads contained SNPs unique to this insertion and displayed a downstream unistrand piRNA profile. RT-PCR and RNA-FISH confirmed that this insertion is transcribed as part of *cactus* mRNAs in follicle cells (Figure S6).

Finally, we investigated the noncoding RNA *CR44619*, which harbors *tirant* insertions in two strains (*A3* and *A7*). Strikingly, these insertions occupy the same position in exon 2, but in opposite orientations: antisense in *A3* and sense in *A7*. RT-PCR confirmed their transcription in the respective strains. Notably, only the antisense insertion led to downstream piRNA production, suggesting that piRNA biogenesis in the cytoplasm is influenced by the sequence content of the transcript (Figure S7). Consistent with the low expression of *CR44619* in follicle cells, piRNA levels from this locus were only moderate.

Together, these findings uncover an unexpected mechanism of transposon silencing where antisense insertions into host gene 3′ UTRs can act as potent unistranded piRNA sources, bypassing the need for canonical piRNA cluster architecture.

### A single 3′ UTR insertion is sufficient to silence *tirant*

To determine whether the *tirant* insertion in *Fs(2)Ket* is necessary and sufficient for piRNA production and silencing, we first generated a hybrid strain in which the second chromosome from *iso-1* was replaced with balancer chromosomes lacking *tirant* piRNA sources. Small RNA sequencing confirmed the loss of piRNA production at the *Fs(2)Ket* locus and a near-complete loss of *tirant*-targeting piRNAs overall (Figure 6A; Figure S8). As expected, this strain exhibited strong *tirant* expression as revealed by RNA-FISH (Figure 6B).

**Figure 6:**
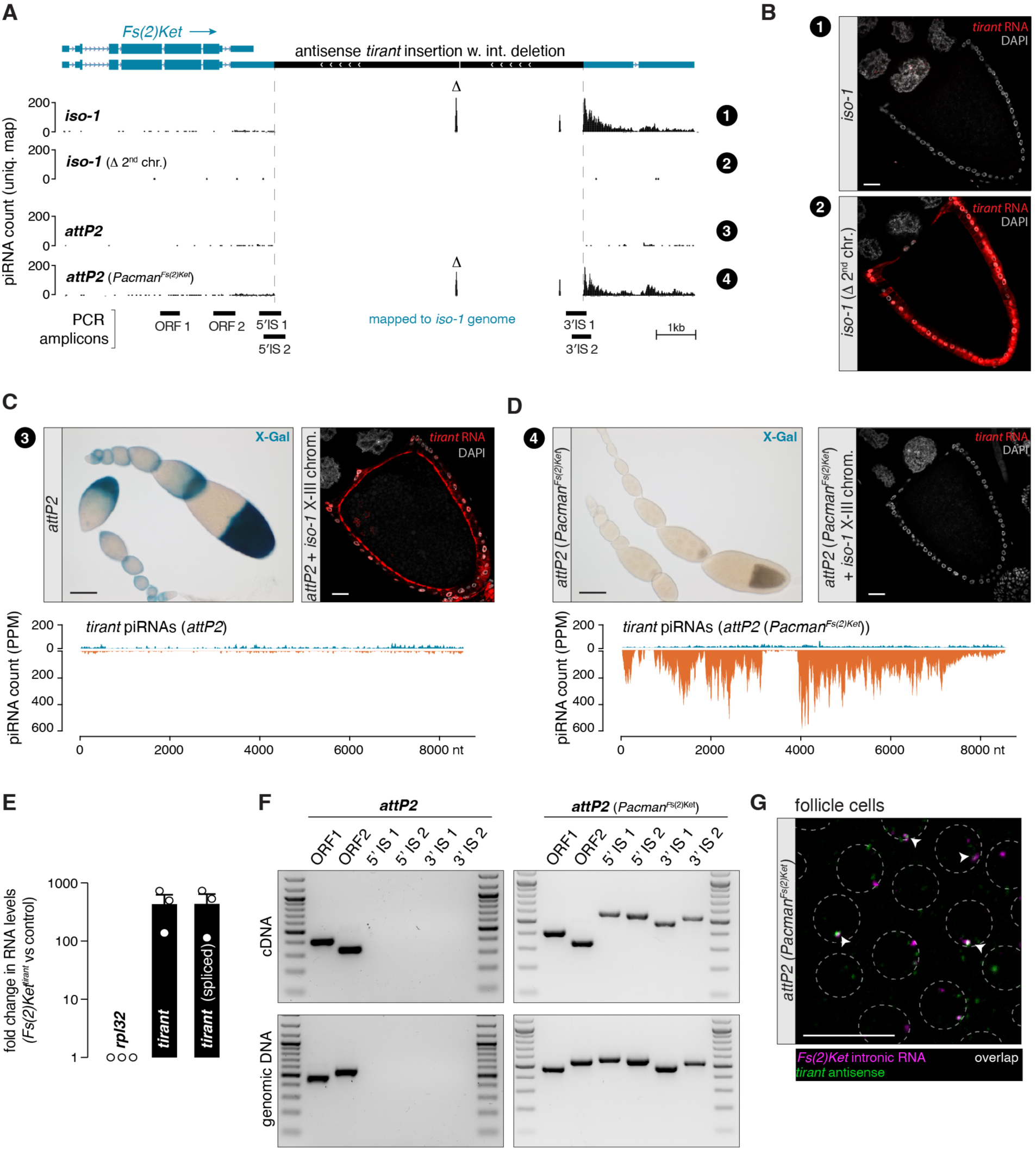
**A single 3′ UTR insertion is sufficient to silence *tirant.*** (A) UCSC browser screenshot of the *Fs(2)Ket* locus in the *iso-1* genome. The four tracks show genome-unique piRNAs (in PPM) from ovaries of the *iso-1* strain, an *iso-1* strain with the second chromosome exchanged by balancer chromosomes (*iso-1(Δ2^nd^ chr.*)), a strain carrying an empty *attP2* landing site, and a strain containing a Pacman clone containing the *Fs(2)Ket* locus from *iso-1* in *attP2* (*attP2 (Pacman^Fs(2)Ket^*)). piRNAs mapping uniquely to the internal deletion of *tirant* are labeled by Δ. Relative genomic positions of analytical PCR amplicons shown in panel D are indicated at the bottom. Scale bar: 1kb. (B) RNA-FISH detecting *tirant* sense transcripts (red) in stage 10A egg chambers from the *iso-1* strain (top) in which *tirant* is silenced and in the *iso-1^Δ^ ^2nd^ ^chr^* strain (bottom), in which *tirant* is expressed. DNA (DAPI) is shown in grey. Scale bar: 20μm. (C) X-Gal staining of an ovariole of the progeny of *tirant* reporter females crossed to males of the empty *attP2* strain (top left), and RNA-FISH detecting *tirant* sense transcripts (red) in a stage 10A egg chamber of a strain carrying the empty *attP2* landing site and the X and 3^rd^ chromosome of the *iso-1* strain (top right). Scale bar: 20μm. Density plot showing ovarian piRNAs mapped to *tirant* (in PPM) isolated from the *attP2* strain (bottom). (D) As in panel C but in the progeny of *tirant* reporter females crossed to males carrying the Pacman clone containing the *Fs(2)Ket* locus from *iso-1* (*Pacman^Fs(2)Ket^*) in the *attP2* landing site. Density plot showing ovarian piRNAs mapped to *tirant* (in PPM) isolated from ovaries carrying the *Pacman^Fs(2)Ket^*clone from *iso-1* in attP2 (bottom). (E) RT-qPCR showing fold change (log10 scale) of *tirant* and spliced *tirant env-F* transcripts between ovarian RNA from the strains used for RNA-FISH in (D) versus (C). (F) RT-PCR for *Fs(2)Ket* on ovarian RNA isolated from the empty *attP2* strain (left), and *attP2 (Pacman^Fs(2)Ket^*) strain (right) and the respective control PCR on genomic DNA. Positions of amplicons indicated in panel A. Marker: 100bp DNA ladder. (G) RNA-FISH detecting *Fs(2)Ket* transcripts (intronic probes; magenta) and antisense *tirant* transcripts (green) in the *attP2 (Pacman^Fs(2)Ket^*) strain. Arrowheads indicate foci with both signals co- localizing. Scale bar: 20μm.

We next asked whether a single genic insertion is sufficient to silence *tirant* in trans. To this end, we reconstituted the *Fs(2)Ket* locus from *iso-1* in a *tirant* naïve strain using a 19 kb genomic BAC (*Pacman^Fs(2)Ket^*) encompassing the entire *Fs(2)Ket* gene and the antisense *tirant* insertion. This construct was integrated into the *attP2* landing site of a strain lacking *tirant* piRNAs (Figure 6C). Small RNA sequencing from *Pacman^Fs(2)Ket^* ovaries revealed robust piRNA production from the *tirant* insertion, including reads spanning the internal deletion (Figure 6A, D). Downstream of the insertion, piRNAs displayed a unistranded profile, similar to the endogenous locus in *iso-1* (Figure 6A).

When crossed to the *tirant-lacZ* reporter strain, the *Pacman^Fs(2)Ket^* transgene silenced reporter expression completely. Moreover, the transgene was sufficient to silence the full set of endogenous *tirant* insertions in an *iso-1* background lacking its native second chromosome, as shown by RNA-FISH and RT-qPCR (Figure 6C-E). RT-PCR and RNA-FISH confirmed transcription of the transgene as a chimeric *Fs(2)Ket– tirant* transcript (Figure 6F, G).

This transgene assay enabled us to directly test whether host gene transcription is required for piRNA production and silencing. We inserted a strong cleavage/polyadenylation signal upstream of the *tirant* insertion in the *Pacman^Fs(2)Ket^* transgene, just downstream of two experimentally validated cleavage/polyadenylation sites of the short *Fs(2)Ket* isoform (*Fs(2)Ket-SV40*, Figure 7A-B). RT-PCR confirmed efficient transcriptional termination upstream of the *tirant* sequence (Figure 7C). Strikingly, flies carrying this modified transgene exhibited a complete loss of *tirant* silencing, as evidenced by their inability to silence both the *tirant* reporter and endogenous *tirant* expression (Figure 7D). Consistent with this, small RNA sequencing revealed a >30-fold reduction in *tirant*-derived piRNAs relative to the original *Pacman^Fs(2)Ket^* strain (Figure 6D, 7E).

**Figure 7:**
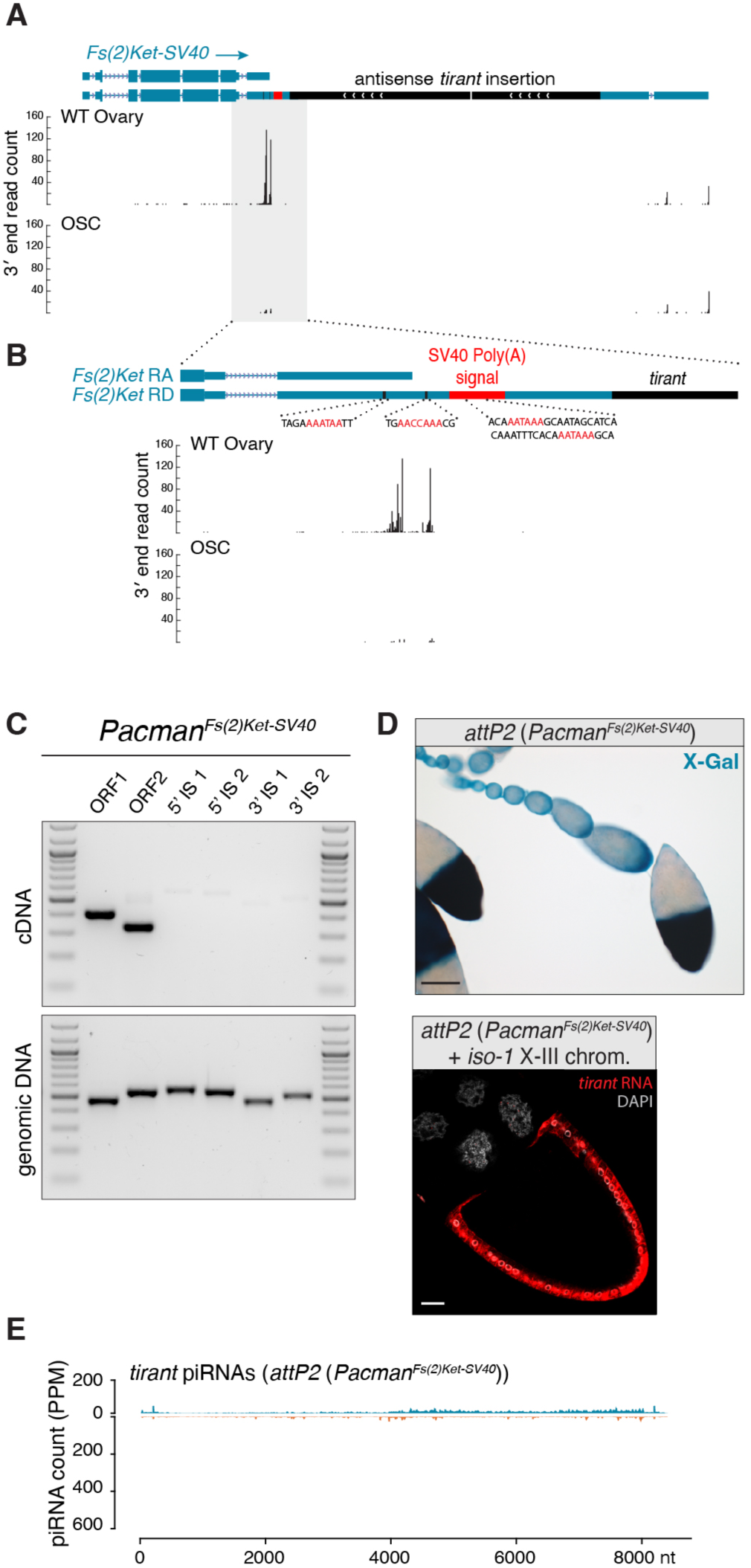
piRNA-mediated silencing of *tirant* requires the *tirant* insertion to be part of the *Fs(2)Ket* transcript. (A) Schematic of the *Fs(2)Ket* locus contained in the Pacman clone in which a strong polyadenylation signal from the SV40 3′ UTR was inserted between the end of the short isoform and the *tirant* insertion (top). Shown below is the 3′ end profile of long-read RNAs from wild type (WT) ovaries (genotype: *tj*-Gal4*>arr2^GD^*) and cultured ovarian somatic cells (OSCs) highlighting 3′ ends of the different *Fs(2)Ket* transcripts. (B) Zoom-in of the part highlighted in grey in panel A, showing predicted polyadenylation signals from the endogenous *Fs(2)Ket* locus and the inserted *SV40* 3′ UTR. (C) RT-PCR from ovarian RNA (top) and PCR from genomic DNA (bottom) from flies harboring the Pacman*^Fs(2)Ket-SV40^*transgene. Marker: 100bp DNA ladder. (D) X-gal staining of ovarioles from the progeny of *tirant-lacZ* reporter females crossed to males carrying the *Pacman^Fs(2)Ket-SV40^* transgene. Scale bar: 100μm (top). RNA-FISH detecting *tirant* sense transcripts (red) in a stage 10B egg chamber of a strain carrying an insertion of the *Pacman^Fs(2)Ket-SV40^*in the *attP2* landing site in a genetic background containing the X and the third chromosomes from the *iso-1* strain. DNA staining (DAPI) shown in grey. Scale bar: 20μm. (E) Density plot showing ovarian piRNAs (in PPM) isolated from the *Pacman^Fs(2)Ket-SV40^* strain and mapped to the *tirant* consensus sequence.

Collectively, these experiments demonstrate that a single antisense *tirant* insertion, when transcribed as part of a host gene 3′ UTR, is sufficient to initiate robust piRNA production and drive trans-silencing of *tirant* in the ovarian soma. Critically, this activity does not rely on *tirant*-intrinsic promoters but instead depends on transcription from the host gene through the insertion.

### Antisense fragments in the 3′ UTR of an ectopic transgene are sufficient to silence *tirant*

Our findings suggest that antisense *tirant* insertions, whether in *flamenco* or host gene 3′ UTRs, initiate silencing by generating unistranded piRNAs from transcripts exported to the cytoplasm where piRNA biogenesis occurs. If this model is correct, the key requirement for piRNA generation should simply be the presence of an antisense transposon fragment within any cytoplasmic transcript, independent of host gene identity or nuclear RNA processing context.

To directly test this model, we engineered two UAS transgenes. Each encodes GFP followed by an SV40 3′ UTR, into which we inserted a 2 kb *tirant* fragment spanning the 5′ UTR and part of *gag*, in either sense or antisense orientation. This design allowed us to control transgene expression using the Gal4- UAS system (Brand & Perrimon, 1993) while eliminating any host gene-specific features.

When expressed in ovarian follicle cells of flies lacking endogenous *tirant* piRNAs but carrying the *tirant-lacZ* reporter, only the antisense construct triggered strong reporter repression, while the sense construct had no effect (Figure 8A,B). To assess whether this effect extended to active elements, we introduced both constructs into the *iso-1* background lacking its second chromosome, and thus lacking the *Fs(2)Ket-tirant* piRNA source. Once again, only the antisense construct suppressed endogenous *tirant* expression (Figure 8A,B).

**Figure 8:**
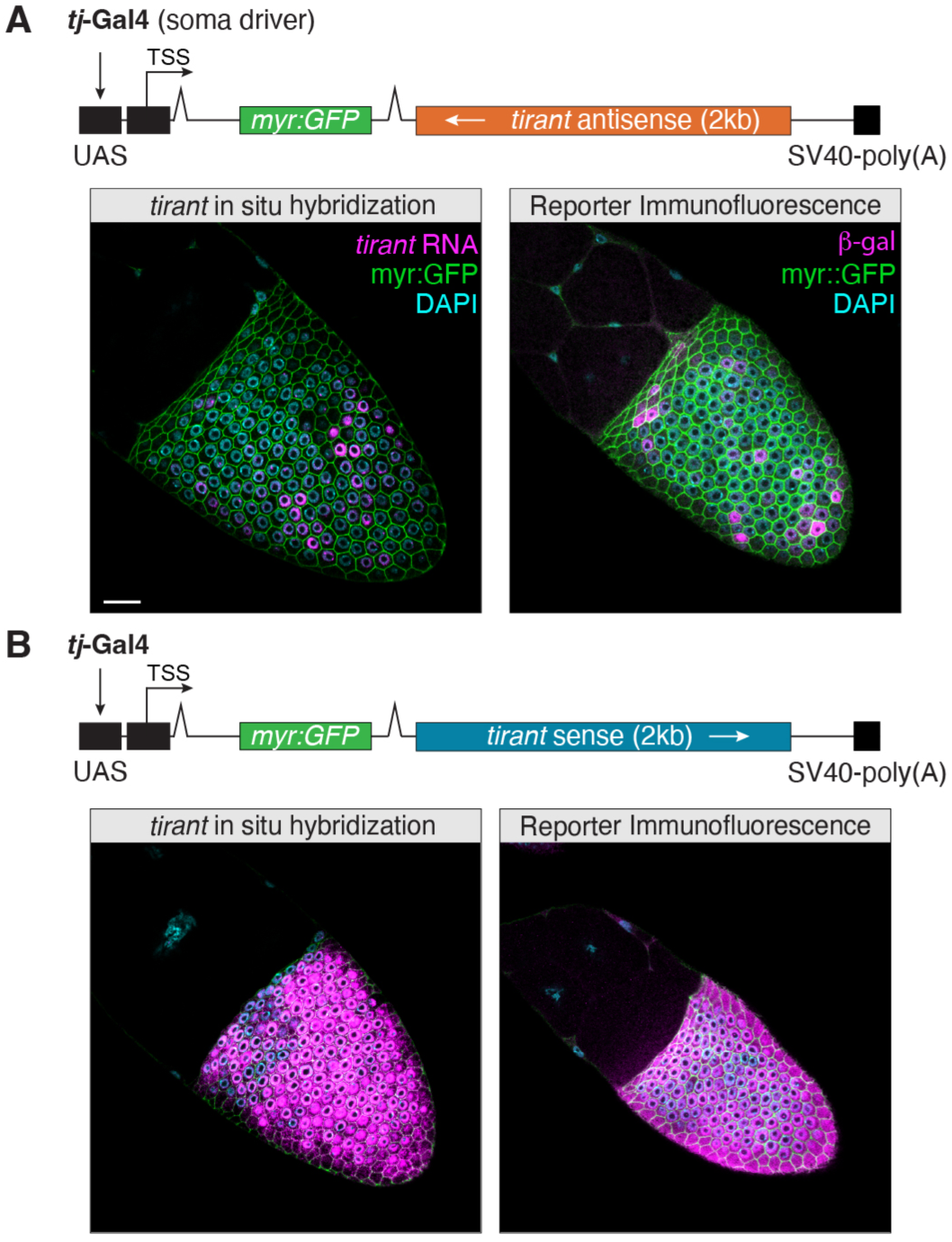
**Antisense fragments in the 3′ UTR of an ectopic transgene are sufficient to silence *tirant.*** (A) Schematic of the artificial *tirant* antisense locus based on a *UAS-myr:GFP* transgene (top). RNA-FISH detecting *tirant* sense transcripts (magenta) in a stage 10B egg chamber in the progeny of a cross between *UAS- myr:GFP*-antisense *tirant* harboring flies and the *tj*-Gal4 driver containing active *tirant* copies (left) or to a *tj*-Gal4 strain containing the *tirant- lacZ* reporter followed by anti-β-Gal staining to detect reporter expression (right). DNA staining (DAPI) in cyan. Scale bar: 20μm. (B) As in panel A but with the *tirant* fragment inserted in the sense orientation. Laser power and gain for the GFP channel in panels A and B was adjusted for similar intensity (GFP levels were considerably lower in the construct with the antisense insertion).

These results demonstrate that an antisense transposon fragment embedded within a cytoplasmic transcript is sufficient for piRNA generation and transposon silencing. Crucially, this process requires neither a canonical piRNA cluster nor a specific host gene context.

## DISCUSSION

Transposable elements are typically neutral or deleterious to host fitness, but in rare instances their insertions can be co-opted for host benefit (Barron *et al*, 2014). In this study, we describe such a case in the recent invasion of the *tirant* retrotransposon into *Drosophila melanogaster*. This horizontally acquired transposable element posed an immediate threat to genome integrity, yet specific insertions into host transcriptional units enabled the emergence of an effective piRNA-based immune response. These findings demonstrate how transposition, the very mechanism fueling transposon spread, becomes the Achilles’ heel of transposons by eventually generating the transcripts required to silence them.

The central conceptual advance of this work is that cytoplasmic transcripts containing antisense transposon fragments are potent and sufficient triggers of piRNA biogenesis in the ovarian soma. This principle holds across three contexts: natural antisense insertions into the *flamenco* piRNA cluster, antisense insertions into host gene 3′ UTRs, and synthetic transcripts bearing antisense fragments. Importantly, these findings demonstrate that piRNA production can occur independently of canonical piRNA-cluster architecture, chromatin context, or host gene identity. Thus, our work broadens the “trap model” of piRNA immunity (Bergman *et al*., 2006; Brennecke *et al*., 2007; Zanni *et al*., 2013): rather than being restricted to specialized clusters, any sufficiently expressed host locus can act as a functional trap for antisense TE insertions, rapidly converting invasive sequences into sources of silencing piRNAs.

The extended trap model provides a highly flexible route to immunity as a single antisense 3′ UTR insertion can initiate piRNA production and confer trans-silencing of active transposons. At the population level, however, the two silencing strategies differ in their evolutionary dynamics. The *flamenco* cluster resides in pericentromeric heterochromatin, a low-recombination environment facilitating the long-term retention and drift-driven fixation of beneficial insertions. In contrast, 3′ UTR insertions occur in euchromatic regions with high recombination rates and may incur fitness costs by interfering with host gene expression. Intriguingly, both *Fs(2)Ket* and *cactus*, the genic piRNA source loci identified here, possess alternative 3′ UTR isoforms, and in each case the *tirant* insertion resides in the less abundant isoform. This suggests a selective balance between achieving silencing and minimizing deleterious effects on gene function. Such constraints may explain the relative rarity of genic piRNA source loci, even though two-thirds of host genes have alternative 3′ UTRs potentially compatible with this strategy (Alfonso-Gonzalez *et al*, 2023; Lee *et al*, 2022). Genic insertions might therefore act as first-line yet transient silencing solutions, eventually giving way to more stable, cluster-based defense mechanisms such as *flamenco*.

The broader significance of the extended trap model is underscored by parallels in other species. In wild koala populations, a single antisense insertion of the KoRV-A retrovirus into a host gene’s 3′ UTR has been linked to piRNA-mediated immunity against this invading virus (Yu *et al*., 2025; Yu *et al*., 2019). Similarly, studies in mammals and mosquitoes have revealed that endogenous retroviral and viral insertions near host gene termini produce antisense transcripts that feed into the piRNA pathway (Konstantinidou *et al*., 2024; Qu *et al*., 2023). Together with our functional data in *Drosophila*, these observations point to a generalizable strategy by which genomes can rapidly evolve defenses against invasive genetic elements, highlighting antisense insertions as critical catalysts of piRNA immunity.

## Limitations of the study

While our findings establish that cytoplasmic antisense transposon RNAs are sufficient to trigger piRNA biogenesis, the molecular mechanisms that selectively channel these transcripts into the pathway remain unclear. In addition, our study focuses on the ovarian soma but several DSPR strains also produce *tirant*-targeting piRNAs in the germline. Although we show that somatic piRNAs are sufficient for silencing, additional roles for germline piRNAs cannot be excluded (Akkouche *et al*, 2013; Yoth *et al*., 2023). Finally, piRNA biogenesis in the germline operates primarily through piRNA-guided slicing and ping-pong amplification. While transcripts with antisense transposon information are also essential in the germline, their licensing into the pathway likely follows distinct rules. Drawing on the recent studies in koalas (Yu *et al*., 2025; Yu *et al*., 2019), sense piRNAs processed from active elements likely trigger antisense piRNA production by cleaving transposon antisense transcripts whose presence relies on the extended trap model presented here.

## ACKNOWLEDGEMENTS

We thank the VBCF and IMBA/IMP/GMI core facilities for support, particularly the NGS facility for sequencing, the BioOptics unit for imaging support, and the VDRC for fly stocks. We thank Peter Duchek and the IMBA Fly & Worm facility for generating transgenic *Drosophila* strains, and Stuart Macdonald, the Drosophila Synthetic Population Resources, and the Bloomington stock centre for sharing fly stocks. Aleksandr Tsarev provided support with sRNA cloning. Members of the Brennecke laboratory and Marianne Yoth gave valuable comments on the manuscript.

## AUTHOR CONTRIBUTIONS

**BR**: Conceptualization; Data curation; Formal analysis; Validation; Investigation; Visualization; Methodology; Writing (original draft); Writing (review and editing).

**LP**: Investigation; Writing (review and editing).

**DH**: Software; Formal analysis; Investigation; Methodology; Writing (review and editing).

**JB**: Conceptualization; Resources; Supervision; Funding acquisition; Visualization; Project administration; Writing (original draft); Writing (review and editing).

**KAS**: Conceptualization; Formal analysis; Investigation; Methodology; Supervision; Funding acquisition; Writing (original draft); Writing (review and editing).

## MATERIALS AND METHODS

### Fly work

All fly stocks used in this study are listed in Table S1. Tissue specific knockdowns were performed as described in (Senti *et al*., 2025). Introgression of *iso-1* chromosomes into the ovarian tissue specific Gal4 strains and RNAi lines was performed by crossing them to *iso-1* flies carrying different combinations of additional balancer chromosomes. To map the silencing capacity of each DSPR strain or the *iso-1* strain to individual chromosomes, we generated different combination of hybrid strains by crossing each DSPR strain to a strain containing balancers on the second and third chromosome, yielding strains that either carry individual strain specific X-chromosomes, autosomal chromosomes or individual autosomal chromosomes.

### Microscopy

**β-gal staining:** Chromogenic β-Galactosidase assays were performed as described in (Handler *et al*., 2013). Briefly, ovaries are dissected into 1X PBS and then fixed for 15 minutes with 0.5% glutaraldehyde in PBS. Samples are rinsed 3 times with 1X PBS and washed once with 1X PBS for 10 minutes. Samples are incubated in staining solution (10mM sodium phosphate buffer, pH 7.0, 1mM MgCl2, 150nM NaCl, 3mM potassium ferricyanide, 3mM, potassium ferrocyanide, 0.1% Triton X-100, 0.05% X-gal) for one hour at 37 degrees. Samples were washed with 1X PBS and mounted on a microscope slide with 75% glycerol. Ovaries were imaged on a Zeiss Imager.Z2 with a Zeiss Axiocam 506 colour camera and a 10x/0.45 plan-apochromat objective.

**HCR fluorescence in situ hybridization:** All RNA-FISH experiments were performed using HCR-FISH (Choi *et al*, 2018) as described in (Luo *et al*, 2020). Samples were mounted with Prolong Diamond and imaged on a Zeiss LSM 880 with a 40x/1.4 EC plan-apochromat Oil DIC or a 63x/1.4 plan-apochromat Oil DIC objectives. Probes were designed as in (Glotzer *et al*, 2022) and are provided in Table S1.

**Immunofluorescence**: Immunofluorescence stainings were performed as described in (ElMaghraby *et al*, 2019) with anti-β-Galactosidase antibodies (Invitrogen A-11132 – 1:1000), and samples were imaged with a Zeiss LSM 880 with a 40x/1.4 EC plan-apochromat Oil DIC objective.

### Molecular work

**Cloning of lacZ reporter and GFP constructs:** The *tirant-lacZ* reporter was cloned as described in (Senti *et al*., 2025). The *tirant-GFP:lacZ* reporter was cloned similarly but using a vector containing a *nlsGFP* fused to the *lacZ* gene. The sequence of the LTR and 5′ UTR of *tirant* were amplified by PCR using a BAC containing an intact copy of *tirant* as a template (BACR48O22) (Hoskins *et al*, 2000). Both reporters were integrated into the *attP40* landing site on the second chromosome (Markstein *et al*, 2008). pJFRC12 (Pfeiffer *et al*, 2012) was used as a backbone to insert the 2kb of *tirant* sequence from the 5′ UTR and beginning of the *gag* gene in the SV40 3′ UTR of the UAS vector. The *tirant* sequences were amplified by PCR from BACR48O22.

**Double short-hairpin construct cloning:** The construct for expressing short hairpin RNAs against *aub* and *ago3* was cloned as described in (Hayashi *et al*, 2016) using sequences given in Table S1.

***Fs(2)Ket* transgenes:** A Pacman containing the whole *Fs(2)Ket* locus from *iso-1* (CH322-190I4) (Venken *et al*, 2009) was injected into *attP2* to generate the *Pacman^Fs(2)Ket^* flies. The *Fs(2)Ket-SV40* Pacman was generated by recombineering as described in (Ejsmont *et al*, 2011). The SV40 sequence was amplified by PCR from the pJFRC12 vector (Pfeiffer *et al*., 2012) and fused to the rpsL cassette via fusion PCR.

**RNA extraction for RT-qPCR and RT-PCR:** Five pairs of ovaries were dissected and homogenized in Trizol. RNA was extracted using Trizol/chloroform followed by isopropanol precipitation and washed once with 80% ethanol. The RNA was treated with DnaseI for 30 minutes and cleaned using a Zymo RRC25 kit following the manufacturer’s instructions. 1ug of RNA was digested a second time with DnaseI and reverse transcribed using the NEB LUNA-RT kit and random primers. qPCR reactions were performed as biological triplicates, with each reaction performed as a technical triplicate. PCR was performed using Taq polymerase, while qPCR used the NEB LUNA qPCR mix (see Table S1 for primer sequences).

**Genomic DNA extraction:** Genomic DNA from roughly 20 adult flies of the same genotype was extracted using the NEB Monarch Spin gDNA Extraction Kit following manufacturer’s instructions. Purified genomic DNA was then used directly for PCR.

**Small RNA-seq:** Small RNAs bound to Argonaute proteins were isolated using TraPR columns as described in (Grentzinger *et al*., 2020) and small RNA-seq libraries from ovaries or dechorionated and washed 0-30 minutes old embryos were generated as described in (Baumgartner *et al*., 2022).

### Computational analyses

**Sequence alignment and phylogenetics analyses:** To place *tirant* within the phylogeny of the *ZAM* subclade of iERVs in *Drosophila melanogaster*, we compared it with previously described consensus sequences of the ZAM subclade members *gypsy5, ZAM, accord,* and *accord2* (Senti *et al*., 2025). We further included two *tirant-*related retroviral consensus sequences, whose fragments are present in the *dm6* reference genome. These were first the initially in *Drosophila yakuba* described iERV *Pifo*, fragments of which are frequently found inserted in *flamenco,* including in *iso-1* (Zanni *et al*., 2013). In addition, we added The *D. simulans* iERV *tirant*S, as partial sequences of this element are harboured in *flamenco* and other heterochromatic sites in *iso-1* (Bargues & Lerat, 2017; Fablet *et al*, 2006). *Pifo* and to a lesser extent *tirant*S represent multiple broken sequences previously described as *tirant*-like in most *Drosophila melanogaster* strains including those lacking active *tirant* (Schwarz *et al*., 2021). For phylogenetical analyses, we annotated the *Pifo*_Dya and *tirant*S_Dsim consensus sequences for open reading frames and aligned the Pol coding region from all iERV consensus sequences in *Drosophila melanogaster dm6* with MACSE2.0 (Ranwez *et al*, 2018), followed by IQ-tree phylogenetic tree estimation (Trifinopoulos *et al*, 2016), and tree display using Dendroscope (Huson & Scornavacca, 2012).

**DSPR genomes:** Long read DNA assemblies were collected from (Chakraborty *et al*., 2019). Annotated genome assembly hubs for the UCSC genome browser were created by aligning reference mRNAs using minimap2 to the published genome assemblies. Repeat annotations were created using RepeatMasker using a curated TE consensus sequence file (attached to the GEO data deposition) (Options: -q -e rmblast

-lib TEconsensus.fa).

***tirant* copy number evaluation:** For the evaluation of the *tirant* copy number in the DSPR genomes, the Illumina DNA sequencing data (King *et al*., 2012) was aligned to the TE-consensus file using bowtie (bowtie --best --strata -f -v 2) and converted to bedgraph files. The median coverage per TE was calculated and normalized to the median coverage of genome unique regions to obtain copy number estimates.

**Analysis of sRNA-seq and RNA-seq:** sRNA-seq was analysed as in (Baumgartner *et al*., 2022). For embryonic samples, piRNA were first normalized to microRNAs and a ratio of ovary/embryo piRNA levels was calculated for piRNAs exclusively expressed in the germline (*burdock*, *F-element*, *DOC*, *I-element*, *mdg3* and *flea*). PingPong scores were calculated as published in (Hayashi, 2022). RNA-seq and long-read RNA-seq were analyzed as described in (Senti *et al*., 2025) and (Voichek *et al*, 2025), respectively.

**Mapping of *tirant* insertions in DSPR genomes:** *tirant* insertions in the different DSPR founder strains were identified by repeat masker (Bao *et al*, 2015) and extracted as individual sequences from each genome. Insertion coordinates were recorded and compiled into Table S1. For insertions of *tirant* that occurred within a gene, the orientation of the insertion relative to the transcription of the gene was also recorded. Production of piRNAs from each insertion was assessed by looking at uniquely mapping sRNA around and inside each insertion.

---

**Figure S1:**
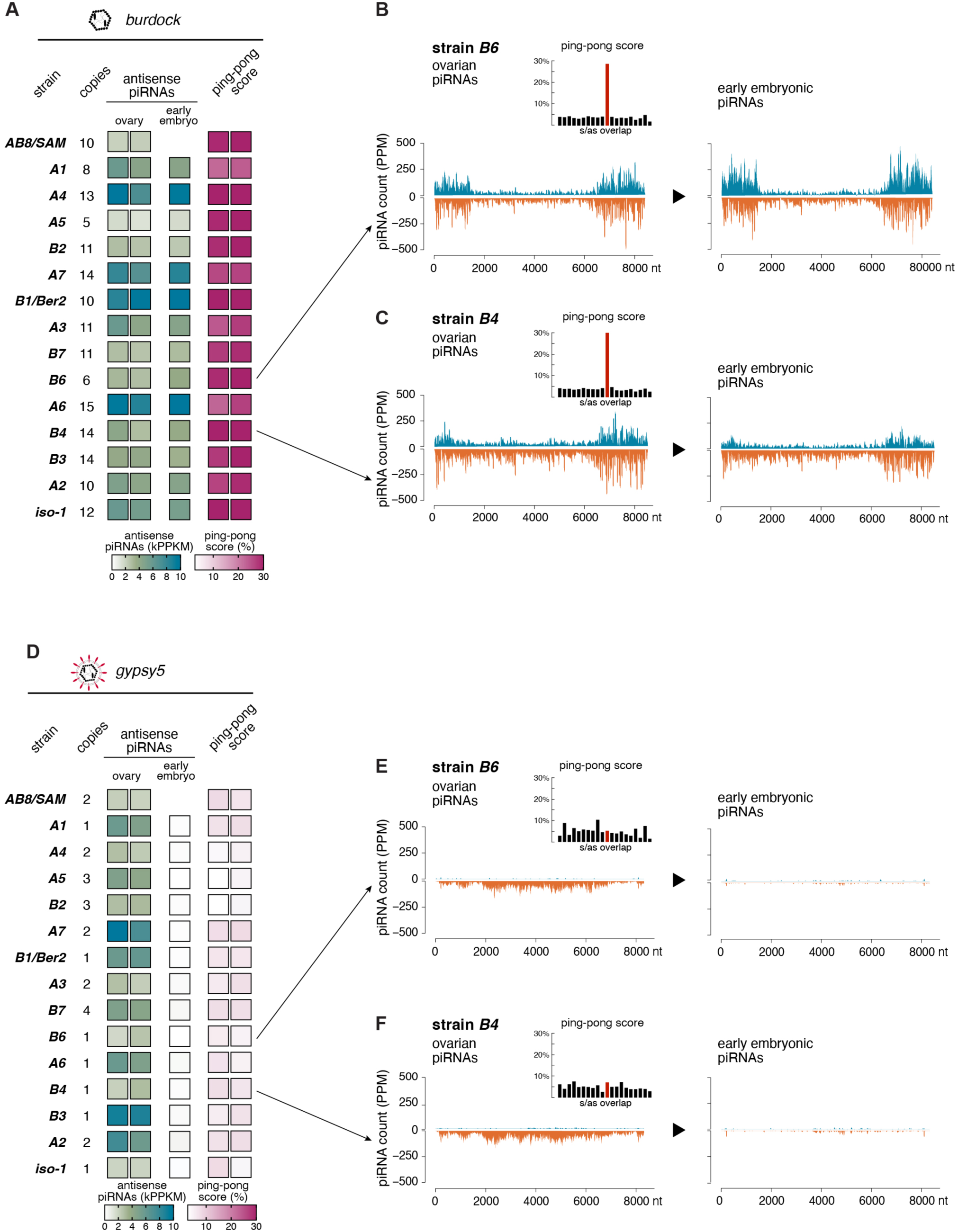
DSPR founder strain piRNAs targeting *burdock* (germline transposon) or *gypsy5* (somatic transposon). (A) Table (left) showing estimated *burdock* copy numbers from genomic Illumina data and heatmaps (right) of 23–29 nt antisense piRNA levels (thousand reads per kb, normalized to 1M miRNAs) mapping perfectly to *tirant* in ovaries and 0–30 min embryos as well as ping-pong scores for ovarian piRNAs. Ovarian small RNAs determined in biological duplicate. (B) Density plots of sense (positive) and antisense (negative) *burdock*-mapping piRNAs (23–29 nt, PPM) along the consensus *burdock* sequence in *B6* ovaries (left) and early embryos (right). Inset: 5′ overlap histogram of ovarian piRNAs (10 nt ping-pong overlap in red). (C) As in (B), for strain *B4*. (D) Table (left) showing estimated *gypsy5* copy numbers from genomic Illumina data and heatmaps (right) of 23–29 nt antisense piRNA levels (thousand reads per kb, normalized to 1M miRNAs) mapping perfectly to *gypsy5* in ovaries and 0–30 min embryos as well as ping-pong scores for ovarian piRNAs. Ovarian small RNAs determined in biological duplicate. (E) Density plots of sense (positive) and antisense (negative) *gypsy5*-mapping piRNAs (23–29 nt, PPM) along the consensus *gypsy5* sequence in *B6* ovaries (left) and early embryos (right). Inset: 5′ overlap histogram of ovarian piRNAs (10 nt ping-pong overlap in red). (F) As in (E), for strain *B4*.

**Figure S2:**
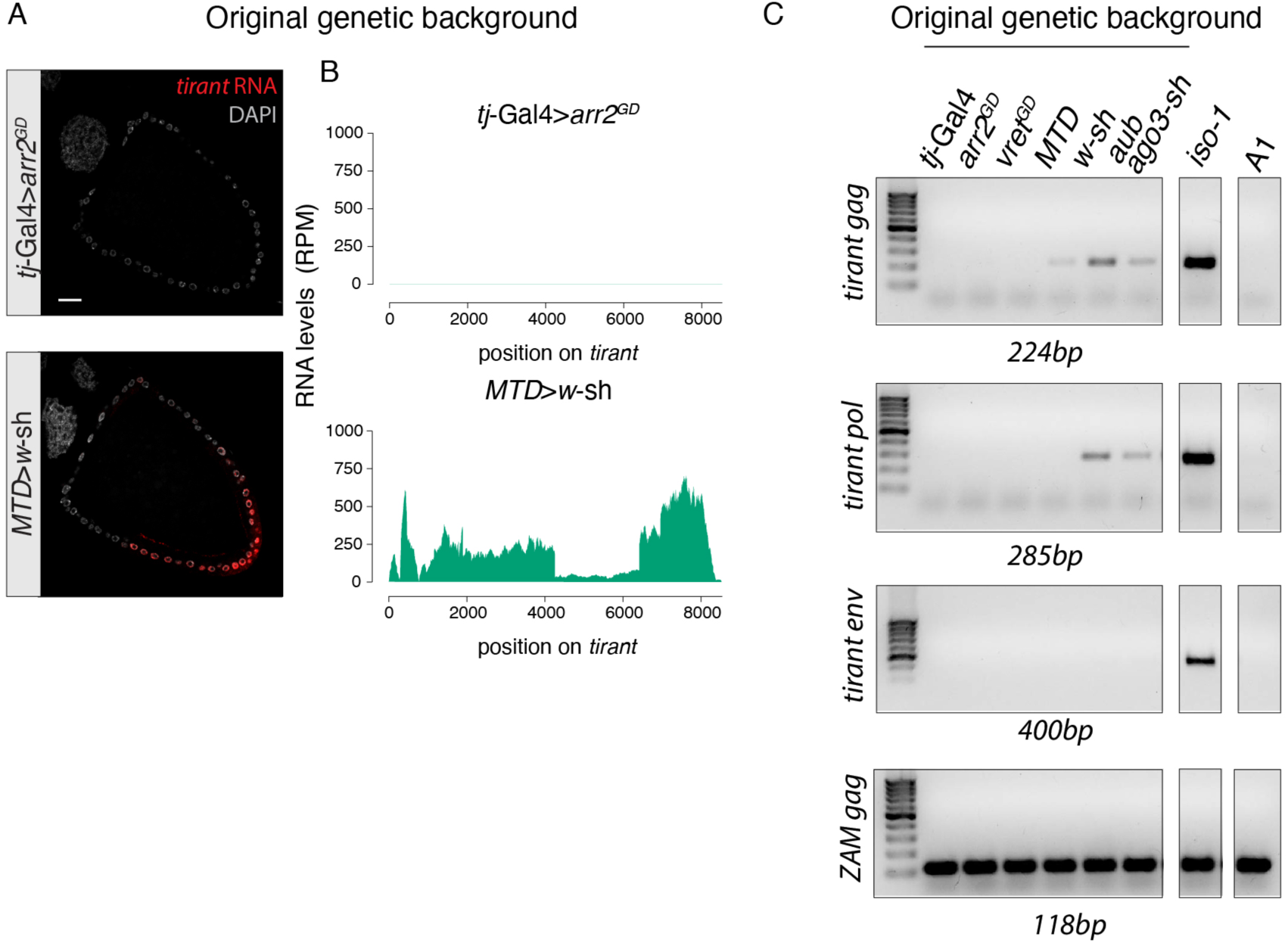
***tirant* is absent or defective in laboratory strains used for piRNA studies.** (A) RNA-FISH detecting *tirant* sense transcripts (red) in ovaries of the original soma (top) and germline (bottom) RNAi strains (control conditions are shown). DNA staining (DAPI) in grey. Scale bar: 20μm. (B) Density of short-read polyA-selected RNA-seq reads (RPM) mapping to the *tirant* consensus sequence from the experimental conditions described in (A) (data from Senti et al, 2025). (C) DNA gel electrophoresis images showing the result of PCR genotyping of genomic DNA of the laboratory strains used in this study to detect presence of *tirant gag*, *pol*, or *env*. DNA from *iso-1* was used as a *tirant*-containing positive control and the *tirant*-free DSPR founder strain *A1* was used as negative template control. The *ZAM* PCR amplicon served as PCR positive control for all strains. DNA marker: 100bp DNA ladder.

**Figure S3:**
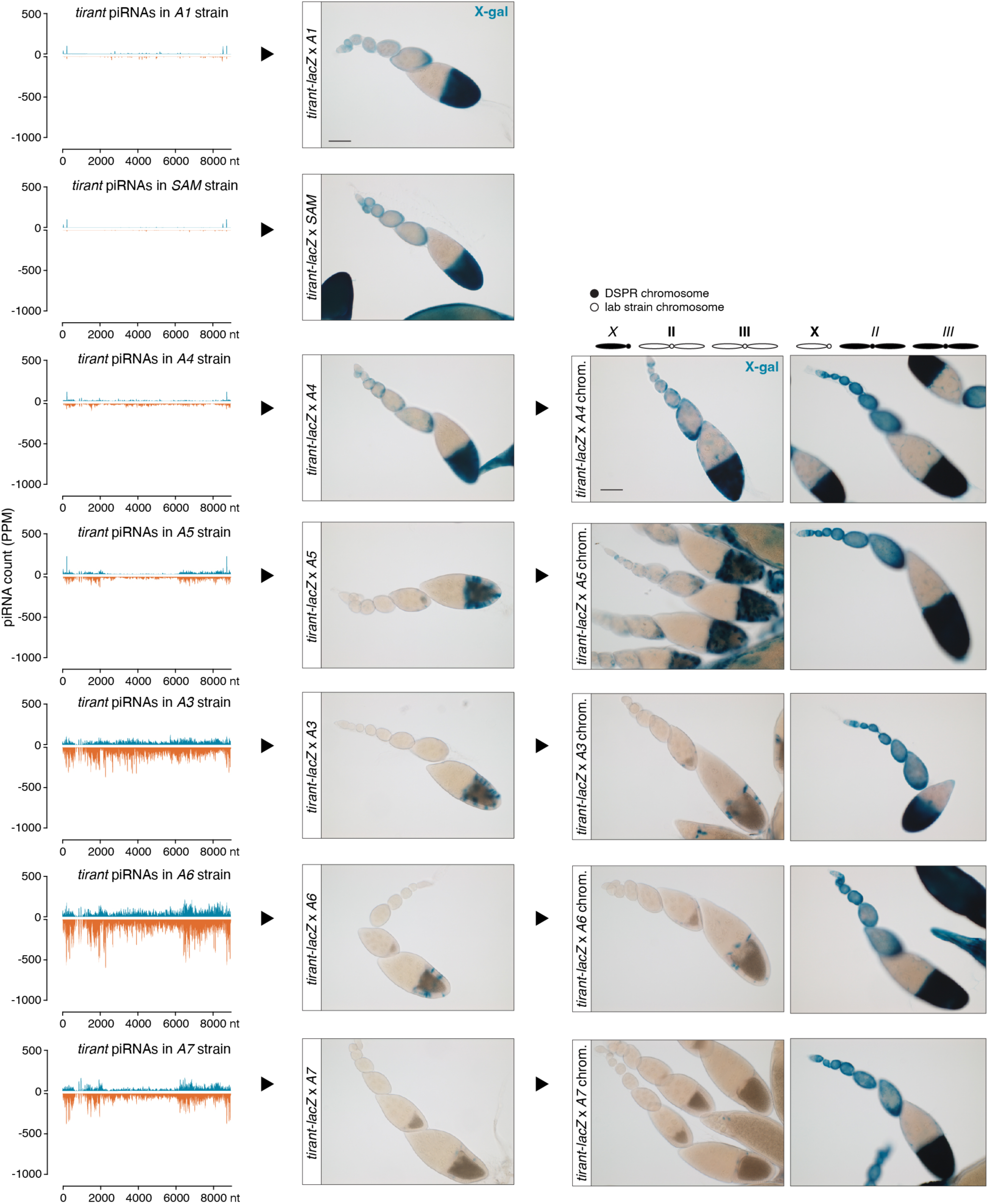
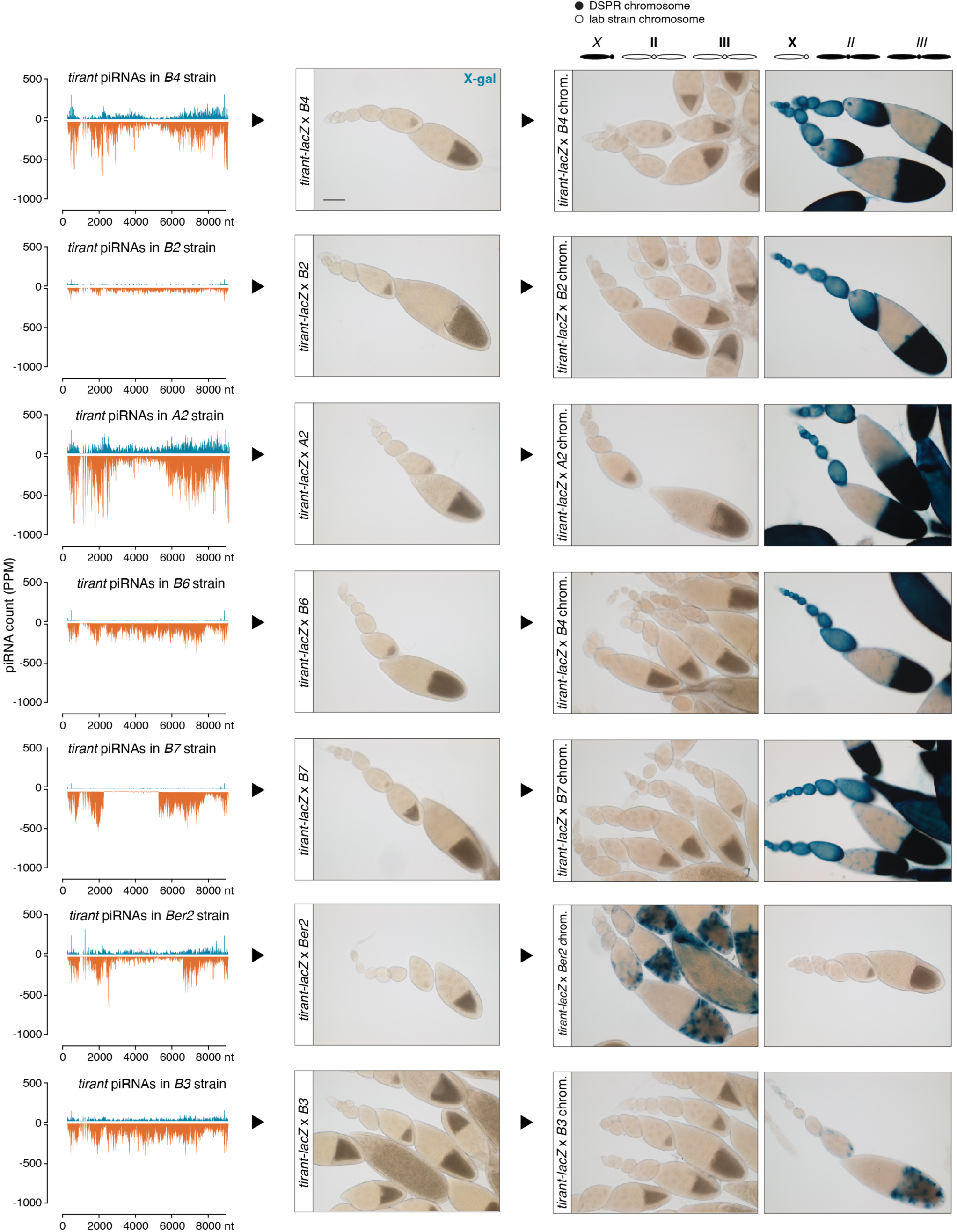
Systematic analysis of the silencing capacity of each DSPR strain and the respective contributions of X chromosome and the pair of autosomes. Shown are the X-gal staining of ovarioles summarized in Figure 3D and the respective profiles of piRNAs mapping to *tirant* in each DSPR strain. piRNA density plots and X-gal stainings from *A1*, *B2*, and *A6* are the same as in Figure 3 and displayed for comparison.

**Figure S4:**
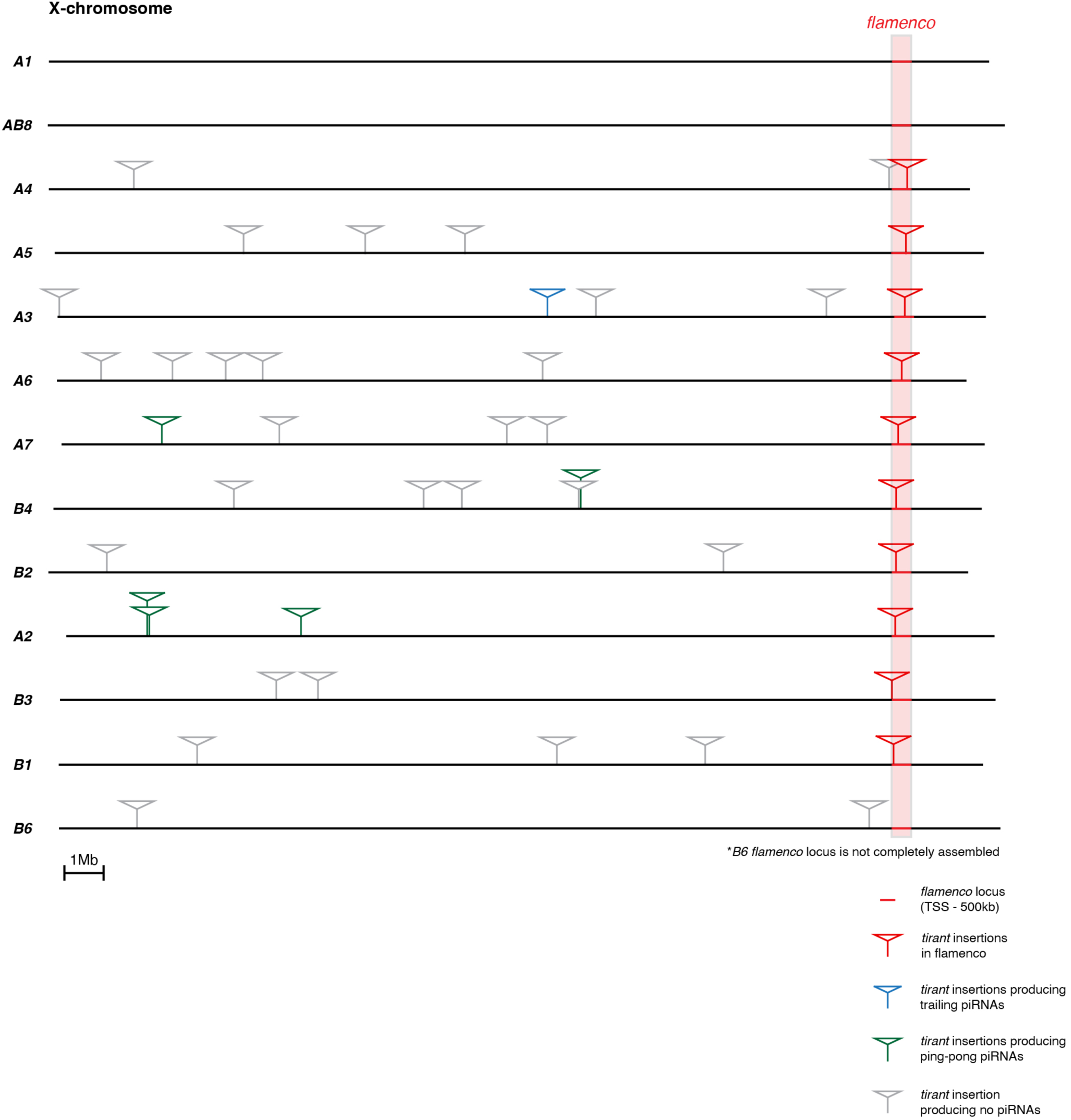
All X chromosomal *tirant* insertions in the DSPR strains. The schematic plot shows the localization of all X chromosomal *tirant* insertion in each DSPR strain based on the respective long read genome assembly (Chakraborty et al, 2019). Patterns of genome-unique piRNAs in each DSPR strain surrounding the respective insertion were visually inspected in UCSC browser tracks to classify insertions into: insertions producing ping-pong piRNAs (green; in strain *A7*, *B4*, and *A2*), insertions producing trailing piRNAs (blue; *A3*), insertions inside *flamenco* (red), insertions with no piRNA pattern (grey). Scale bar: 1Mb.

**Figure S5:**
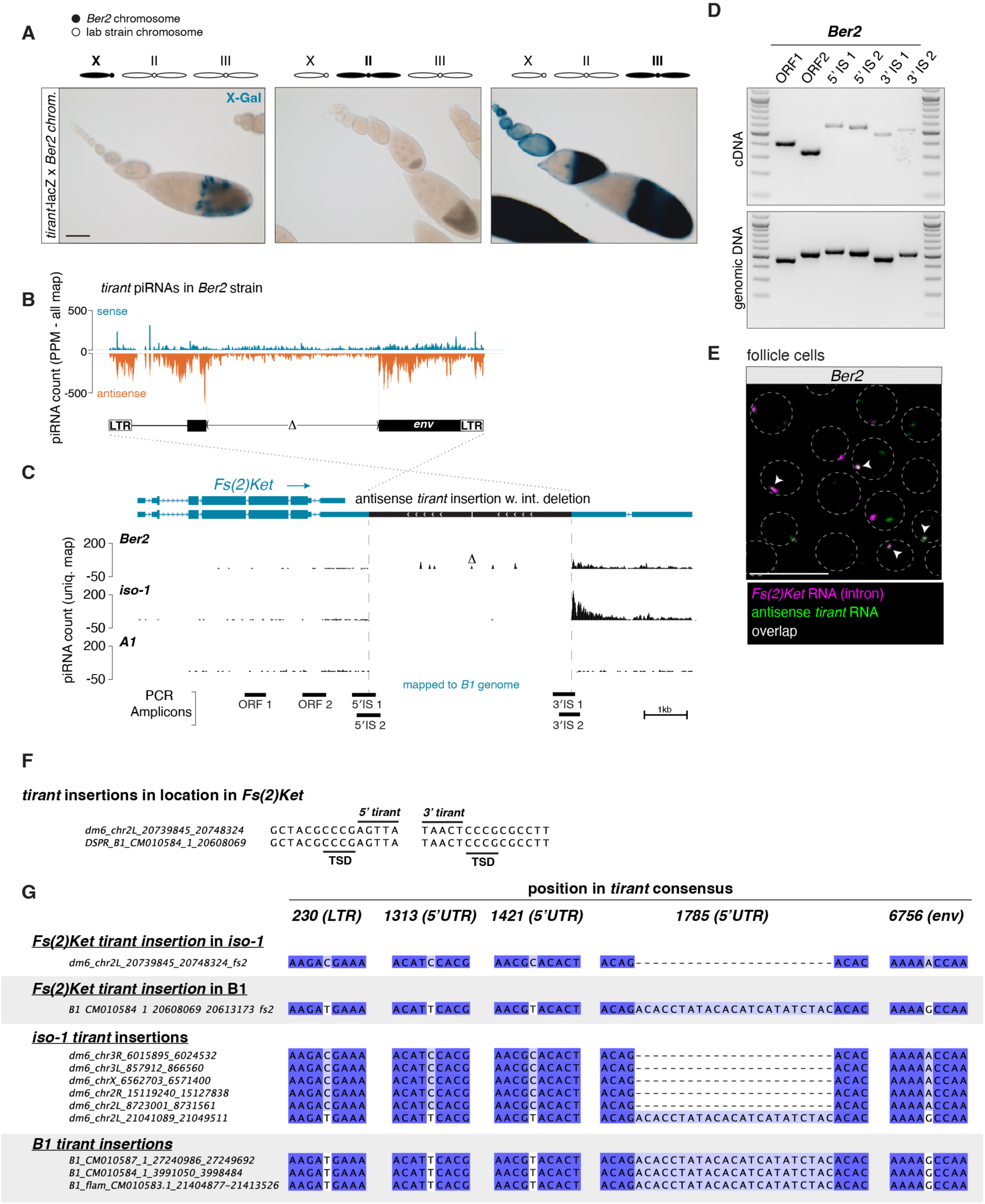
A *tirant* insertion in the *Fs(2)Ket* 3′ UTR is sufficient to produce piRNAs in the *B1*/*Ber2* strain. (A) X-gal stainings of ovarioles from the progeny of crosses between transgenic *tirant-lacZ* reporter females and males harboring either the X (left), 2^nd^ (middle), or 3^rd^ chromosome (right) from the *Ber2* strain as indicated. Scale bar: 100μm. Density plot of piRNAs (PPM) mapping to the *tirant* consensus sequence isolated from *Ber2* ovaries. Dashed lines mark the region with distinctly lower piRNA coverage, corresponding to the deletion in the *tirant* insertion in *Fs(2)Ket*. UCSC browser screenshot of the *Fs(2)Ket* locus in the *B1* genome. The three tracks show genome-unique piRNAs (in PPM) that were sequenced from ovaries of *Ber2* (top), *iso-1* (middle), and *A1* (bottom), each mapped to the *B1* long read genome assembly. piRNAs mapping uniquely to the internal deletion of *tirant* are labeled by Δ. Relative genomic positions of analytic PCR amplicons shown in panel D are indicated at the bottom. Scale bar: 1kb. (D) RT-PCR on RNA and PCR on genomic DNA from the *Ber2* strain to detect chimeric transcripts between *Fs(2)Ket* and *tirant*. Positions of PCR amplicons are shown in panel C. Marker: 100bp DNA ladder. (E) RNA-FISH detecting *Fs(2)Ket* (intronic probes; magenta) and antisense *tirant* (green) transcripts in stage 8 egg chambers of the *Ber2* strain. Arrowheads indicate foci with both signals co-localizing. Scale bar: 20μm. (F) Insertion site sequences of *tirant* copies in *Fs(2)Ket* in the *iso- 1* and *B1* strains. The target site duplications (TSD) flanking the *tirant* LTRs from both 5′ and 3′ ends are indicated. (G) Sequence alignment showing informative parts of the *tirant* insertions in the *Fs(2)Ket* 3′ UTR present in *iso-1* or *B1* together with a subset of other *tirant* insertions in these strains. Shown are all SNPs and an insertion in the 5′ UTR that differ between the insertions in *iso-1* and *B1*.

**Figure S6:**
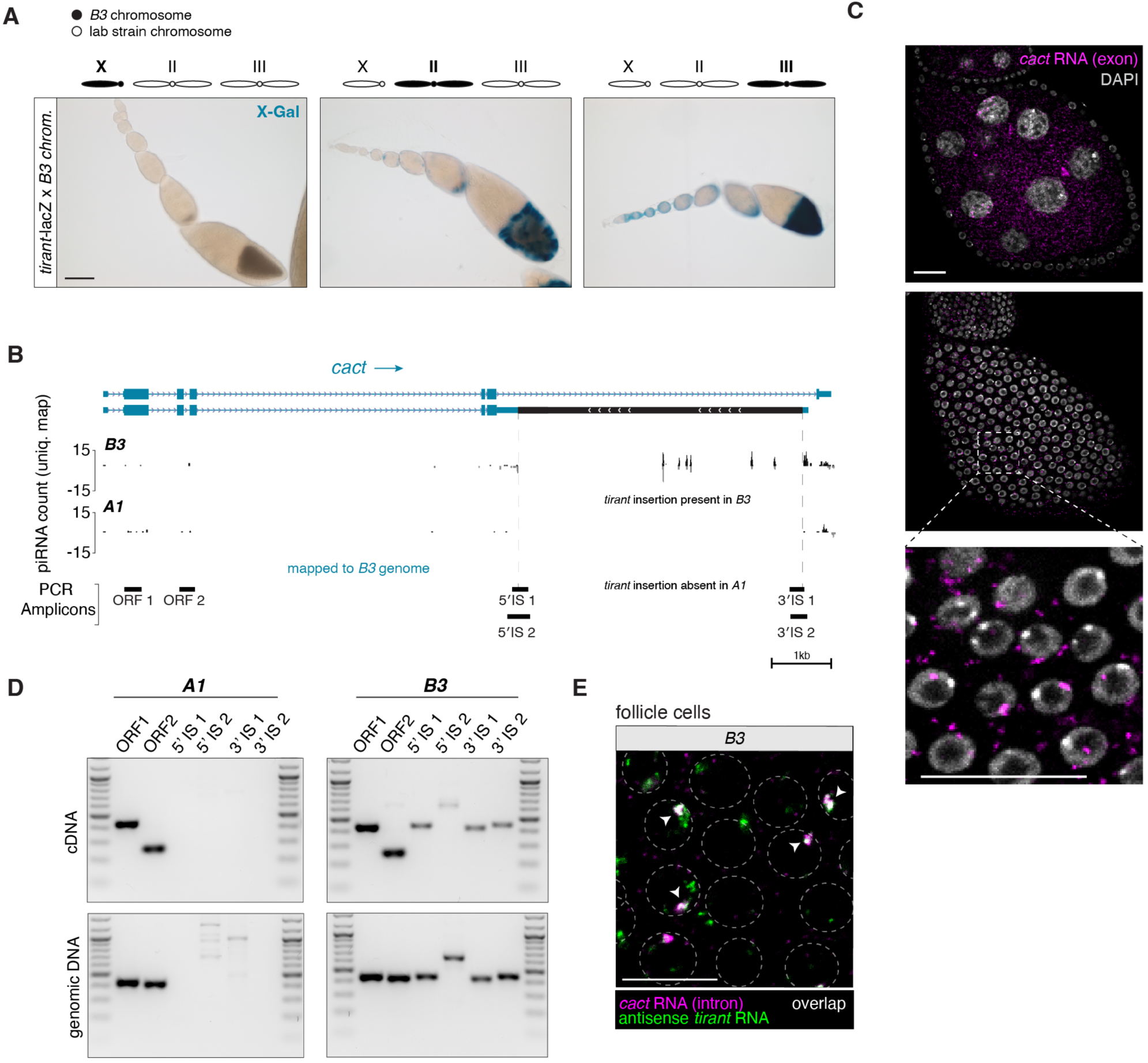
A *tirant* insertion in the *cactus* 3′ UTR is sufficient to produce piRNAs in the *B3* strain. (A) X-gal stainings of ovarioles from the progeny of crosses between transgenic *tirant-lacZ* reporter females and males harboring either the X (left), 2^nd^ (middle), or 3^rd^ chromosome (right) from the *B3* strain. Scale bar: 100μm. (B) Genome-unique piRNAs mapping to the *cact* locus in the *B3* genome showing the two annotated *cact* isoforms. piRNAs from *A1* are shown as negative control. Genomic positions of analytical PCR amplicons shown in panel D are indicated at the bottom. Scale bar: 1kb. (C) RNA-FISH detecting *cact* transcripts (exonic probes) in the germline (top) or somatic cells (middle) of *A1* ovaries with zoom-in from the boxed part shown below. Scale bar: 20μm. (D) RT-PCR on cDNA and PCR on genomic DNA from the *A1* and *B3* strains to detect chimeric transcripts between *cact* and *tirant*. Positions of PCR amplicons are shown in panel C. Marker: 100bp DNA ladder. (E) RNA-FISH detecting *cact* (intronic probes; magenta) and antisense *tirant* (green) transcripts in follicle cells of ovaries from the *B3* strain. Arrowheads indicate foci with both signals co-localizing. Scale bar: 20μm.

**Figure S7:**
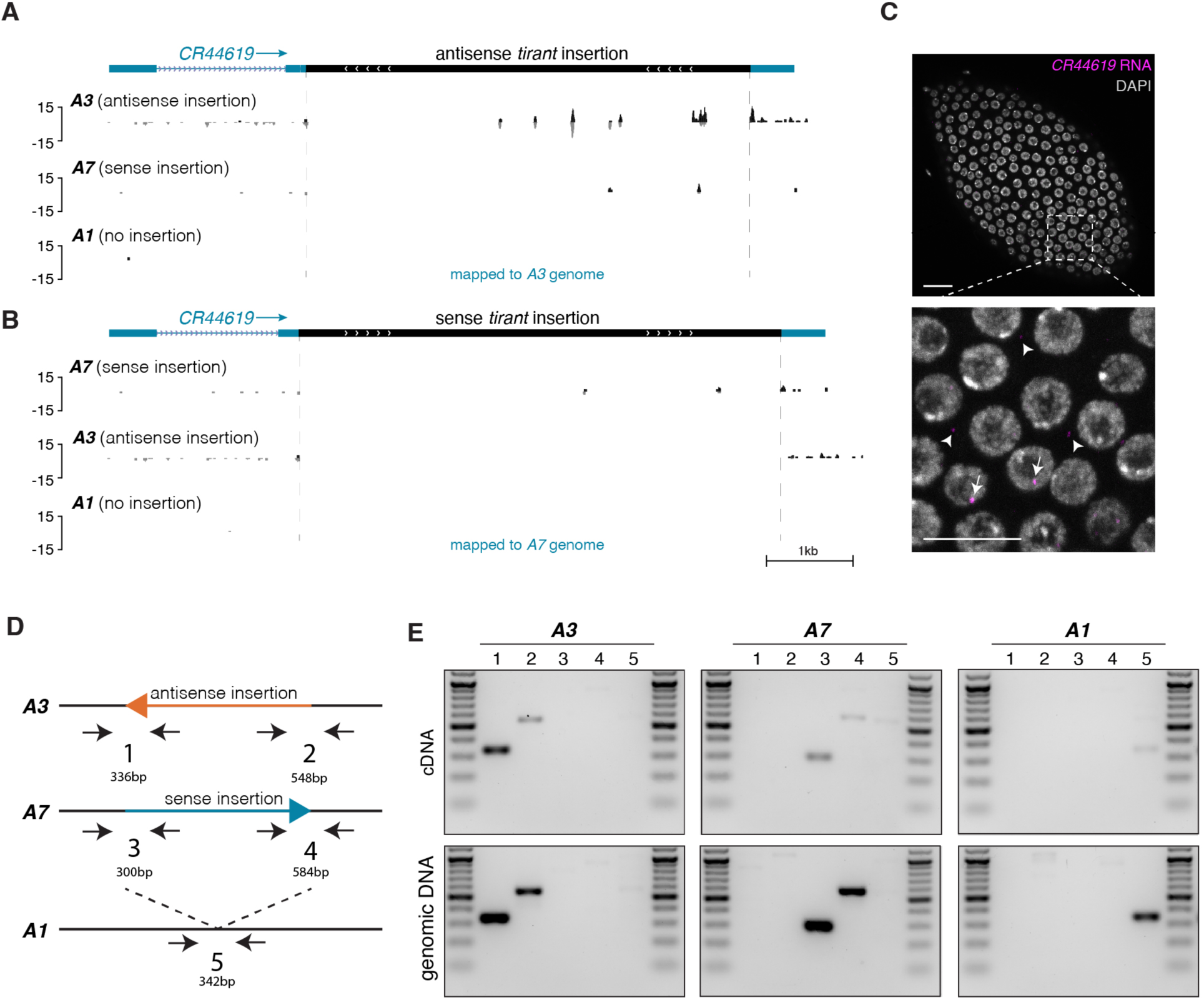
A *tirant* insertion in the long non-coding RNA *CR44619* produces low amount of piRNAs, exclusively in the antisense orientation. (A) UCSC browser screenshot of the *CR44619* locus in the *A3* genome. The three tracks show genome-unique piRNAs (in PPM) from ovaries of the *A3* (top), *A7* (middle), or *A1* strain (bottom). (B) As in panel A but with genome-unique piRNAs mapped to the *A7* genome. (C) Maximum intensity projection of RNA-FISH signal detecting *CR44619* transcripts (magenta) in a stage 8 egg chamber. The image below is a zoom-in of the boxed part in the top panel. Arrows indicate *CR44619* signal in the nucleus and arrowheads signal in the cytoplasm. DNA (DAPI) is shown in grey. Scale bar: 20μm. (D) Schematic showing the positions of PCR amplicons and their expected sizes used in panel E to test the presence of the *tirant* sequence as part of the *CR44619* RNA in the *A3*, *A7*, and *A1* strains. (E) RT-PCR on cDNA and PCR on genomic DNA from *A1* and *B3* to detect chimeric transcripts between *CR44619* and *tirant*. Positions of PCR amplicons are shown in D. Marker: 100bp DNA ladder.

**Figure S8:**
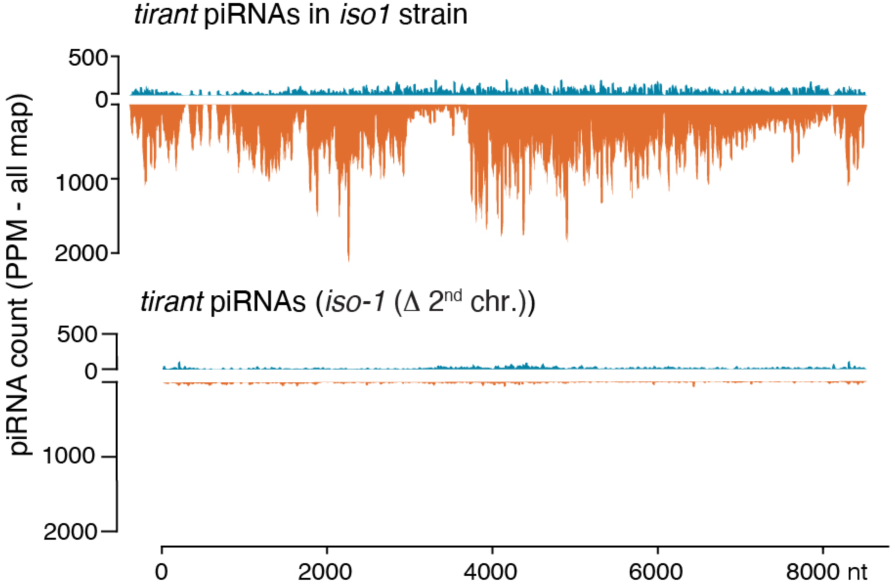
**The *iso-1^Δ^ ^2nd^ ^chr^* strain does not produce piRNAs against *tirant.*** Density plot showing piRNAs (in PPM) mapping to the *tirant* consensus sequence from ovaries of the *iso-1* strain (top) and from the *iso-1(Δ2^nd^ chr.*) strain (bottom). The top plot is reused from Figure 5B for comparison.

